# The locus of recognition memory signals in human cortex depends on the complexity of the memory representations

**DOI:** 10.1101/2021.06.15.448519

**Authors:** D. Merika W. Sanders, Rosemary A. Cowell

## Abstract

According to a “Swiss Army Knife” model of the brain, cognitive functions such as episodic memory and face perception map onto distinct neural substrates. In contrast, representational accounts propose that each brain region is best explained not by which specialized function it performs, but by the type of information it represents with its neural firing. In a functional MRI study, we asked whether the neural signals supporting recognition memory fall mandatorily within the medial temporal lobes (MTL), traditionally thought the seat of declarative memory, or whether these signals shift within cortex according to the content of the memory. Participants studied objects and scenes that were unique conjunctions of pre-defined visual features. Next, we tested recognition memory in a task that required mnemonic discrimination of both simple features and complex conjunctions. Feature memory signals were strongest in posterior visual regions, declining with anterior progression towards the MTL, while conjunction memory signals followed the opposite pattern. Moreover, feature memory signals correlated with feature memory discrimination performance most strongly in posterior visual regions, whereas conjunction memory signals correlated with conjunction memory discrimination most strongly in anterior sites. Thus, recognition memory signals shifted with changes in memory content, in line with representational accounts.

---

The Swiss Army knife analogy of brain function (Cosmides and Tooby 1994) asserts that the human mind contains many functionally distinct cognitive “tools” – e.g., for language acquisition, reinforcement learning, visual perception – each employing different mechanisms that emerged as independent adaptive specializations. This analogy has greatly influenced cognitive neuroscience, encouraging researchers to focus on the separability of cognitive functions and map them to distinct neuroanatomical substrates (e.g., Aggleton and Brown 1999; Kanwisher 2006; Duncan et al. 2019). Applied to memory, the Swiss Army knife analogy manifests as the “Multiple Memory Systems” view, under which theories of memory posit multiple separable systems for different modes of learning and storage, underpinned by distinct neural substrates (Sherry and Schacter 1987; Tulving 1987; Tulving and Schacter 1990; Nadel 1992). More recently, some theories derived from this general framework have rejected one-to-one mappings between cognitive functions and focal brain structures, instead mapping mnemonic functions to distributed networks (Ranganath and Ritchey 2012; Rugg and Vilberg 2013; Thakral et al. 2017; Hebscher and Voss 2020). However, an important legacy of the Swiss Army knife view is that most research questions still center on a cognitive function, such as episodic memory retrieval, and ask which brain region, distributed network, or neural mechanism supports it. That is, cognition is still carved into subdomains that correspond to functional “tools”, and these tools still serve as the cognitive constructs whose (putatively fixed) neural substrates must be identified (Cowell et al. 2019).

A partial departure from this approach has been made by theories that invoke information content to explain the roles of different brain regions in memory (e.g., Davachi 2006; Diana et al. 2007; Shimamura 2011). But many such accounts still explain brain regions’ contributions via a *combination* of function and content, e.g., claiming the hippocampus (HC) underpins recollection (a mnemonic function) of contextual details (a type of content; Davachi et al. 2003; Eichenbaum et al. 2007; Staresina et al. 2011; Staresina et al. 2013; Bastin et al. 2019). We and others have suggested that “content” is more than just a label to be used *in combination with* mnemonic functions or processes, such as recollection; instead, content may be the primary principle driving how memory and perception map onto the brain at the systems level (Bussey and Saksida 2002; Graham et al. 2010; Nadel and Peterson 2013; Cowell et al. 2019).

One theory that departs fully from the Multiple Memory Systems approach is the representational-hierarchical account of perception and memory (Bussey and Saksida 2007; Cowell et al. 2010). This account proposes that the role of each brain region within the ventral visual-medial temporal lobe (MTL) pathway is best explained not by which cognitive function it performs, but by the type of information it represents. Building on well-established evidence for a hierarchy of increasingly complex stimulus representations in visual cortex (Hubel and Wiesel 1965; Felleman and Van Essen 1991; Kobatake and Tanaka 1994; Malach et al. 1995; Kanwisher et al. 1997; Kamitani and Tong 2005; Henriksson et al. 2008; Kriegeskorte et al. 2008; Ostwald et al. 2008; Brouwer and Heeger 2009; Drucker and Aguirre 2009; Serences, Saproo, et al. 2009; Cowell et al. 2017), this account suggests that the representational hierarchy extends beyond visual cortex into the MTL. Thus, the pathway houses a graded continuum of representations, beginning with simple visual features such as oriented lines in V1 and culminating in complex multi-modal conjunctions corresponding to episodes or events in HC. Under this account, any region in the pathway should contribute to any perceptual or mnemonic function – e.g., visual discrimination, mnemonic encoding, or retrieval from declarative memory (including via familiarity signaling or pattern completion-like recall) – if it contains neural representations that are useful for the task.

One prediction of this account is that the MTL contributes to perception of complex stimuli such as 3-dimensional objects and complex spatial scenes – this contradicts the “Multiple Memory Systems” notion that the MTL specializes in long-term, declarative memory (Squire and Zola-Morgan 1991). Much empirical work now supports this prediction (e.g., Buckley et al. 2001; Lee et al. 2005; Bartko et al. 2007; Lee et al. 2012). Indeed, the multi-faceted functional contributions of the MTL are now known to extend to statistical learning (e.g., Schapiro et al. 2014), decision-making (e.g., Palombo et al. 2015), and future simulation (e.g., Thakral et al. 2020) – notably, almost always for tasks involving complex or associative stimuli.

Thus, a key plank of the evidence for a one-to-one mapping between long-term, declarative memory and the MTL has been eroded: we know that the MTL performs non-mnemonic functions. The complementary plank of evidence for this one-to-one mapping concerns whether long-term, declarative memory can be supported by structures outside of the MTL. The representational-hierarchical account suggests that it can, but this prediction has not been thoroughly tested. Some recent studies suggest support for it: early visual regions underlie recognition memory for orientation (Cooke et al. 2015) and spatial location (Thakral et al. 2013; Karanian and Slotnick 2018); pattern completion-based recollection occurs outside of HC for stimuli that do not contain rapidly-learned associations (Ross et al. 2018; Gardette et al. 2022). But, to our knowledge, no study has examined whether recognition memory signals can be made to shift location within the brain by manipulating the complexity of the representations that must be recognized.

We tested this prediction with an fMRI study of recognition memory in humans. We constructed visual objects and scenes defined by unique conjunctions of precisely specified features. This allowed us to manipulate the complexity (i.e., simple, feature-level versus complex, conjunction-level) of the representations that need to be retrieved for recognition. We used objects and scenes with the expectation that both stimulus sets would engage a range of brain regions along the ventral visual-MTL pathway, with the possibility that the transition from feature to conjunction representations would occur slightly earlier in the pathway for objects than for scenes (e.g., Lee et al. 2012; Erez et al. 2016; Cowell et al. 2017). Our key manipulation was the contrast between feature memory and conjunction memory, for both stimulus sets. During recognition memory retrieval in the scanner, we measured feature-based and conjunction-based mnemonic discrimination performance, along with blood oxygenation level-dependent (BOLD) signatures of feature-based and conjunction-based memory. We predicted, following the representational-hierarchical account, that manipulating the complexity of mnemonic information would shift the location of memory signals within the ventral visual-MTL pathway, for both stimulus sets, from posterior visual regions for feature memory to anterior regions for conjunction memory. Further, we predicted that posterior, feature-based memory signals would be most predictive of feature-based mnemonic discrimination performance, whereas anterior, conjunction-based memory signals would best predict the mnemonic discrimination of conjunctions.

## Materials and Methods

### Design Overview

We used novel objects and novel, everyday scenes, with each item composed of the conjunction of three binary features (see Figures 1 and 2, and the *Stimuli* section). We tested each participant on two separate days, one for each stimulus set (objects, scenes). The order of stimulus sets across the two days was counterbalanced across participants. Participants first entered the scanner and viewed a stream of stimuli (fMRI data from this “study phase” are not presented in this report). Participants exited the scanner for a short break, then returned to the scanner for a recognition memory task involving the stimuli they had just viewed in the study phase. For both objects and scenes, test stimuli were either Novel, Recombination, or Familiar. In Novel stimuli, all individual features and conjunctions of features were entirely novel. In Familiar stimuli, both the individual features and the conjunction of those three features had been seen in the study phase. In Recombination stimuli, the individual features were familiar from having been studied as parts of other items, but the conjunction of those three features was novel.

**Figure 1.**
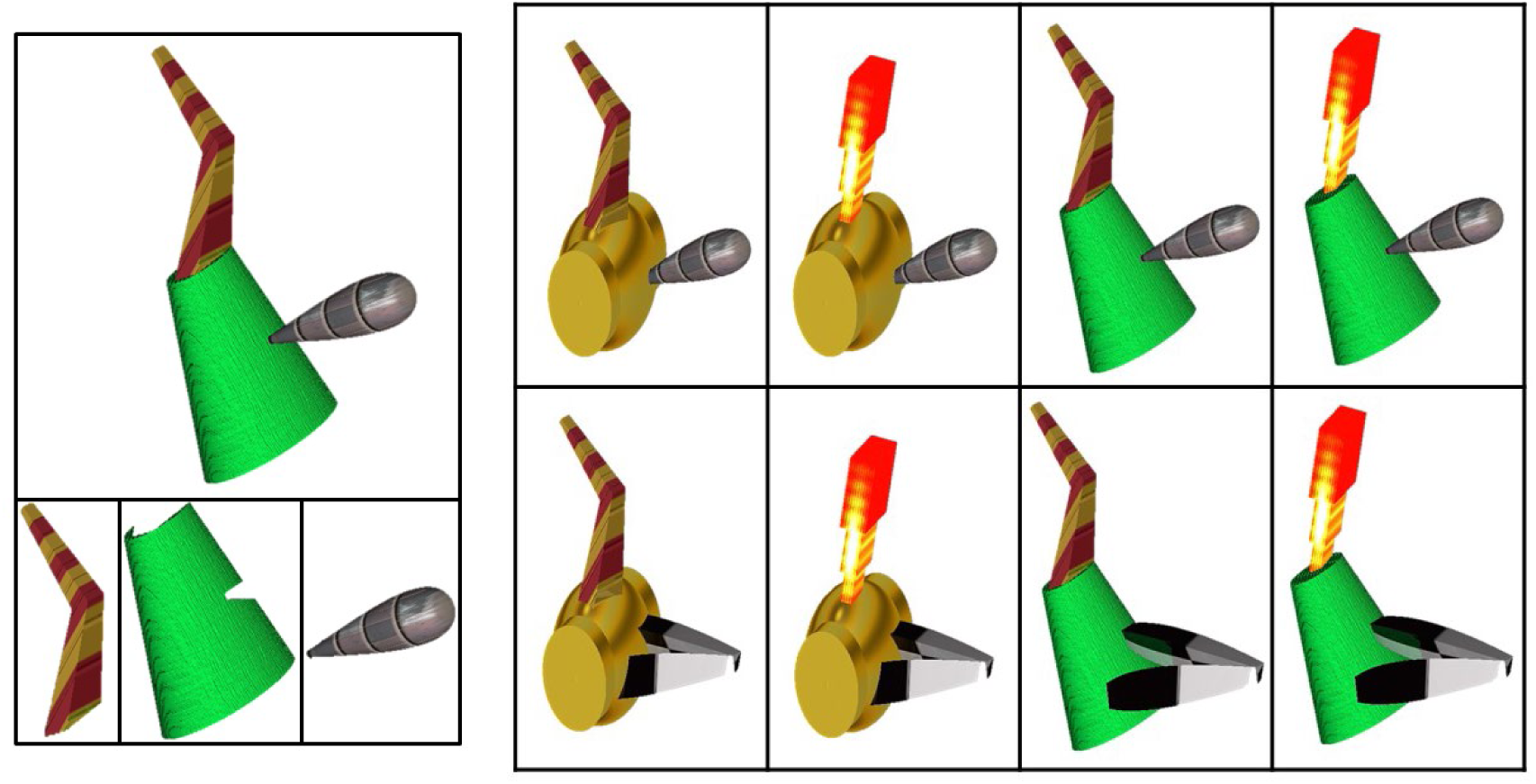
Fribble Stimulus Set Examples. *Left*: a whole, conjunctive Fribble (top) and its tail, body and head features (bottom, left to right); *Right*: A family containing eight unique Fribbles, constructed from three binary features (tail, body and head).

**Figure 2.**
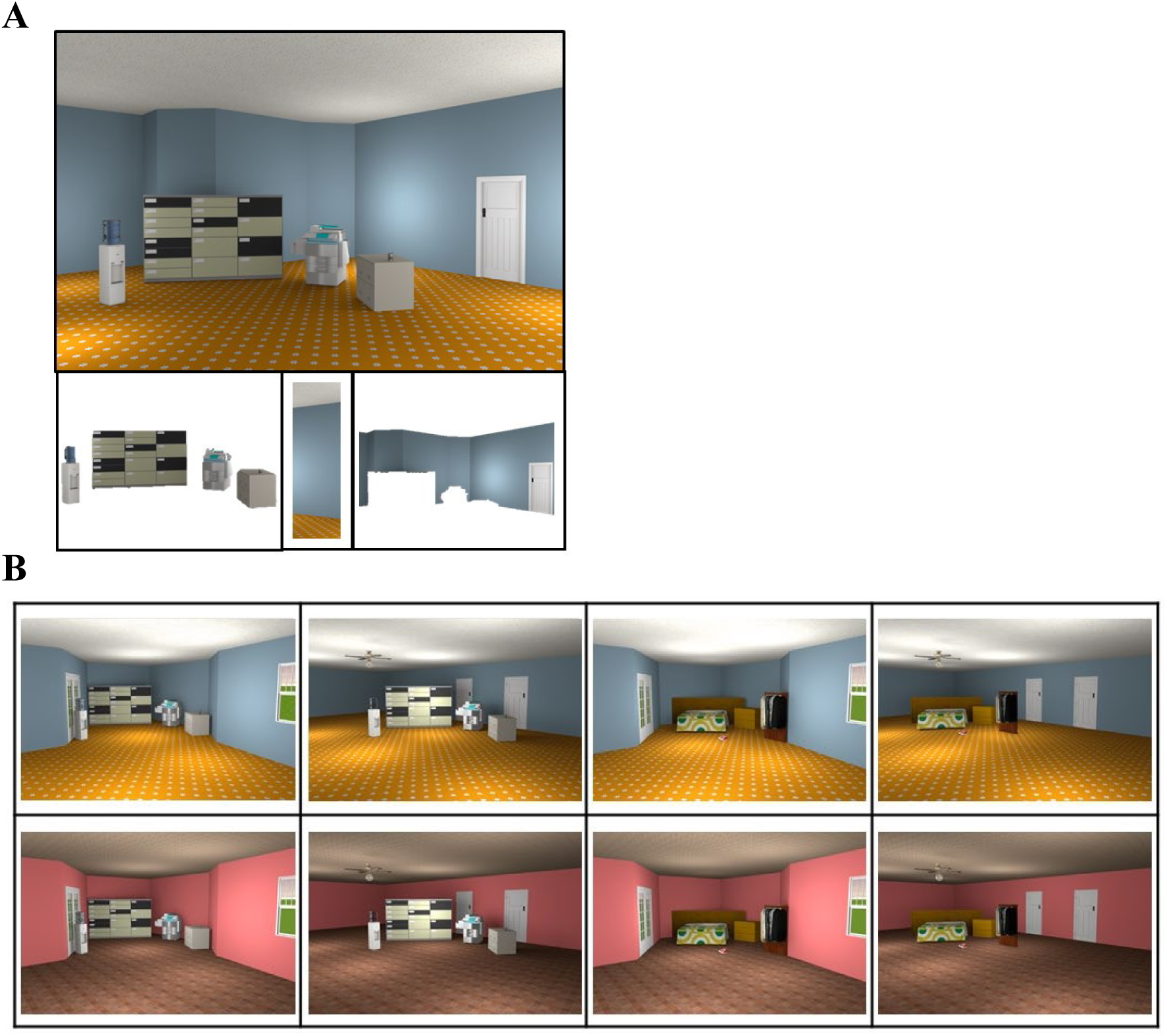
Scene Stimulus Set Examples. *A*: A conjunctive Scene (top) and its features, furniture, color and room shape (bottom, left to right); *B*: A family containing eight unique conjunctive Scenes composed using three binary Scene features.

We assessed feature memory and conjunction memory – in terms of both behavioral performance and neural signals – by comparing different pairs of memory trial types. To assess behavior, we used *d’*, calculated from hits and false alarms using Signal Detection Theory, to measure mnemonic discriminability. To assess neural memory signals, we used a BOLD signal contrast. We assessed *feature memory* by measuring *d’* and the BOLD contrast for Novel versus Recombination trials. This is because the novelty/familiarity of individual features is the only mnemonic factor discriminating these two trial types (both Novel and Recombination stimuli possess a novel conjunction, but only Novel stimuli possess novel features). We assessed *conjunction memory* via *d’* and the BOLD contrast for Recombination versus Familiar trials. Here, the novelty/familiarity of the conjunction of features is the only mnemonic factor discriminating the two trial types (both Recombination and Familiar possess familiar features, but only Recombination stimuli possess a novel conjunction). This approach tests feature memory divorced from conjunction memory, and conjunction memory divorced from feature memory, which delivers finer control of the content of mnemonic representations than, for example, earlier work demonstrating cortical reinstatement in early visual areas during retrieval of visual information (e.g., Thakral et al. 2013).

#### Participants

Twenty-three participants were recruited from the University of Massachusetts Amherst community and gave their written informed consent. All participants spoke English fluently, had normal or corrected-to-normal vision, no history of neurological illness, and no contraindications for MRI scanning. Participants were compensated $25/hour with an additional performance-based bonus up to $10 per scan session.

#### Stimuli

Two distinct stimulus sets were created using graphical software: novel 3-D objects and scenes. The novel 3-D objects, called Fribbles, were created using Strata Design 3D CX 7.5 (Williams 1998; Barry et al. 2014). Each Fribble was a unique conjunction of three features (3-D components referred to as “tail”, “body”, and “head”) and belonged to a “family” (Figure 1). Within a family, there were two possible variants for each feature and those variants were unique to that family. Therefore, a family contained eight unique Fribbles corresponding to all possible conjunctions of that family’s binary features. We created four Fribble families in total.

Novel scenes were created using Sweet Home 3D, an indoor planning software. Each Scene was a unique conjunction of three features (“room shape”, “color”, and “furniture”) and belonged to a “family” (Figure 2). Within a family, there were two possible variants for each feature and those variants were unique to that family. Consequently, each family comprised eight unique Scenes and we created four Scene families.

#### Task

During the study phase, participants completed a 1-back repetition detection task. Some inter-stimulus intervals (ISIs) between study trials contained null trials, during which participants saw a white central fixation cross that dimmed briefly, either once or twice, and participants could press either response key whenever it occurred. No response to 1-back repetitions was required on the first trial of a run, or on trials immediately following a null. Before entering the scanner, participants practiced this task on a laptop computer by completing a series of 32 trials (including 3 immediate repeats) with stimuli that were subsequently never seen during the experiment.

During the break between study and test, participants received instructions on how to distinguish between the three mnemonic stimulus classes (Familiar, Recombination, and Novel). During test, participants indicated the class of the stimulus on the screen using three button box keys: (1) “Familiar” (studied in the previous session); (2) “Recombination” (made of features studied in the previous session, combined in a new way); or (3) “Novel” (not studied in the previous session in any form). In the test phase, null trials occurred during every ISI.

For both study and test, participants were instructed to respond as accurately as possible while the stimulus was still on the screen and were informed that a performance-based bonus was available.

#### Experimental Design

Each participant completed ten study scans, or “runs” (fMRI data not presented here), and six test scans (Figure 3). Stimuli were displayed on a 32” LCD monitor positioned at the head end of the magnet bore, which participants viewed via a mirror on the head coil.

**Figure 3.**
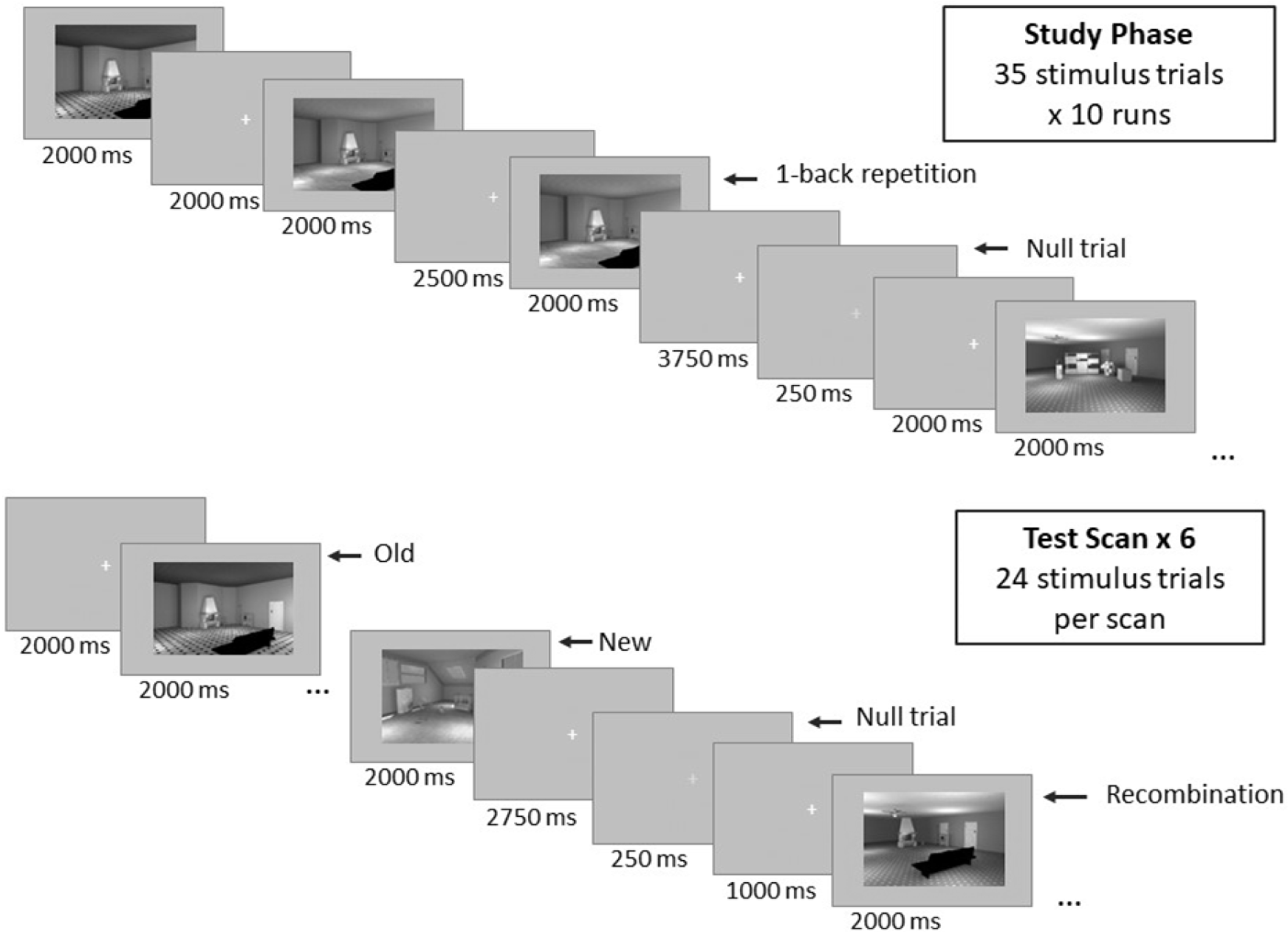
Experimental Design. During the study phase, participants indicated if the current image was identical to the previous image (i.e., 1-back repetition detection task). During the test scan, participants discriminated between Familiar, Novel, and Recombination stimuli. Both the study phase and test scans featured null trials in which participants indicated when a fixation cross dimmed. Images are presented here in grayscale but appeared in color for participants.

In the study phase, stimulus presentation order was blocked by family within each run (e.g., stimuli from one Familiar family were presented in even-numbered study runs, and from the other Familiar family in odd-numbered runs), but stimuli within a run were presented in random order. Within each study run there were 35 stimulus trials: four repetitions each of eight stimuli from a Familiar family, presented singly on a gray background, plus three items repeated a fifth time to create “immediate repeats” for the 1-back task. Each stimulus was presented for 2000ms, immediately followed by an ISI of between 2000ms and 8000ms, during which a white central fixation cross appeared on a gray background. The response window for stimulus trials began 500ms after stimulus onset and extended 1000ms into the ISI.

This design resulted in the presentation of each stimulus at least 20 times for study (4 repetitions of all family members per run, 5 runs per family). This was done in part to increase memory strength (e.g., Atkinson and Juola 1973; Ratcliff et al. 1985; Ratcliff et al. 1990; Jacoby et al. 1998) and in part to increase the precision of BOLD estimates (although neuroimaging data from the study phase are not presented here). As many as 20 repetitions have been used in prior studies of declarative memory, where they were found to dramatically enhance learning and retention (Walsh et al. 2023).

At least 500ms after the end of the response window, for 15 of the 35 ISIs in each study run, a null trial occurred. Null trials occurred only during ISIs of between 3000ms and 8000ms in length. During a null trial, the white cross dimmed to gray for 250ms, either once (for ISIs under 6000ms) or twice (for ISIs 6000 – 8000ms). We randomized the exact onset of the dimming event(s) within the ISI, with constraints to keep dimming events separate and sufficiently distant from the start or end of the ISI. Null trials measured participants’ attention and provided gaps in the sequence with minimal “mind-wandering” (Stark and Squire 2001). After completion of 10 study runs, participants exited the scanner and took a self-paced break before re-entering for the six test runs.

During test runs, 48 Novel, 48 Recombination and 48 Familiar stimulus trials were presented, with 16 unique stimuli from each mnemonic stimulus class shown three times each. Participants performed the same memory task (response options: Familiar, Recombination, or Novel) for every presentation of a test stimulus, including test-phase repeats. The question always asked about the item’s novelty with respect to the study phase, i.e., a “Novel” response meant the item had never been seen in the study phase, although it may have been seen once or twice in the test phase. (Given the large number of study repetitions, familiarity acquired from the study phase always greatly exceeded familiarity acquired from the test phase). Novel and Recombination stimuli were created by combining two features of one family (unstudied for Novel or studied for Recombination) with a third feature of the other unstudied/studied family (Figure 4). Implementing all possible “1 + 2” feature combinations yielded 48 unique stimuli for both Novel and Recombination stimulus classes. However, to match the Familiar stimuli, of which there were only 16 unique exemplars, only 16 out of the 48 possible stimuli were presented for each of Novel and Recombination classes. The subset of 16 stimuli was counterbalanced across participants.

**Figure 4.**
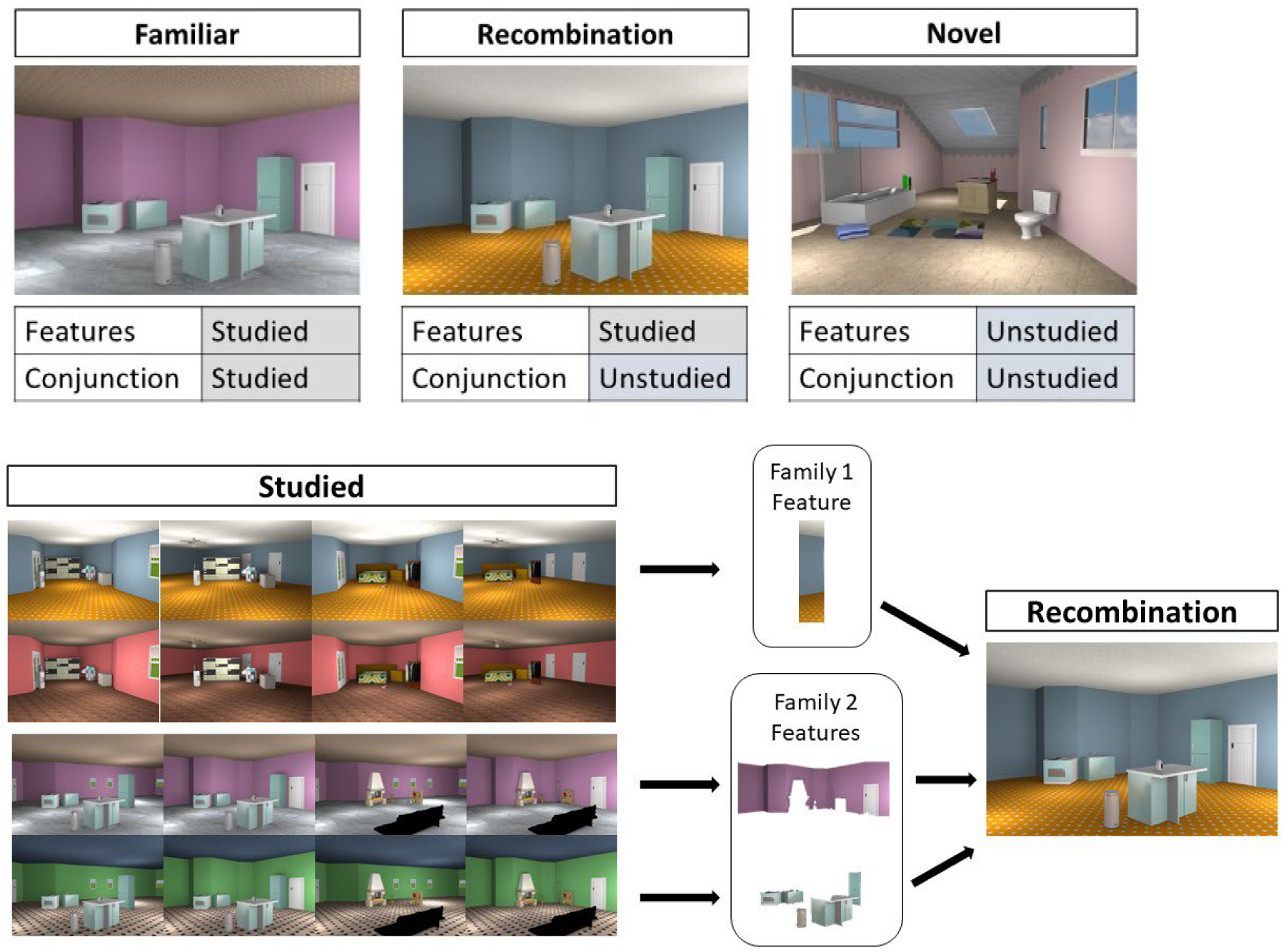
Scene Stimulus Set Recombination Example. *Top*: the three mnemonic stimulus classes differed according to whether features and the conjunction of features were studied in the study phase; *Bottom*: counterbalanced across participants, two of the Scene families were designated to be studied (i.e., presented in the study phase). In this example, a Recombination stimulus is created by combining the color feature from the first family with the room shape and furniture features from the second family. All potential 1 + 2 feature combinations within a given pair of families yielded 48 Recombination stimuli. A Fribble recombination example is shown in Figure S1.

Trial order and spacing of the three stimulus types was determined using the easy-optimize-x MATLAB tool, with the constraints that the three stimulus types appeared equally across the six test runs (i.e., eight of each type per run) and repetition was blocked (i.e., all 48 stimuli were shown once before any stimulus was presented a second time). Each test stimulus appeared for 2500ms, with a randomly assigned ISI between 4000ms and 12000ms. The response window extended 1000ms into the ISI. During every ISI, the null trial commenced only after the response window had elapsed (3500ms after stimulus onset).

#### Image Acquisition

Scanning was performed on a Siemens 3T Skyra scanner equipped with a 64-channel head coil at the University of Massachusetts-Amherst’s Human Magnetic Resonance Center. Functional images were acquired using a T2-weighted EP2D-BOLD sequence (TR: 1250ms; TE: 33ms; flip angle: 70°; FOV: 210mm; 2.5mm^3^ voxels) and forty-two axial slices were acquired during each functional scan. Whole-brain anatomical images were acquired in the middle of the study functional scan session using a T1-weighted (T1w) MP-RAGE sequence (208 sagittal slices; TR: 2000ms; TE: 2.13ms; flip angle: 9°; 1mm^3^ voxels). Functional scans included 20 study runs, 12 test scans and two localizer runs per participant; we used only the data from test runs in the present report.

### Data Analyses

#### fMRI Data Preprocessing

After fMRIPrep’s preprocessing pipeline (see Supplementary Materials for details), functional data was high-pass filtered with a 128s cutoff and functional data from test scan sessions underwent smoothing with a 5mm FWHM Gaussian kernel. The data were analyzed with custom MATLAB scripts using SPM 12 software.

#### Definition of Regions of Interest (ROI)

Despite collecting localizer scans, we ultimately used probabilistic atlases to define ventral visual stream areas (i.e., V1, V2, V3, V3AB, hV4, ventral occipital cortex [VOC], lateral occipital cortex [LOC], temporal cortex

; Wang et al. 2015) and MTL areas (i.e., perirhinal cortex [PRC], parahippocampal cortex [PHC], HC; Ritchey et al. 2015; https://identifiers.org/neurovault.collection:3731). Probabilistic definitions were adequate, or perhaps even preferable, given our theoretical framework: our hypothesis predicted a continuum of memory signals ranging from feature-based in more posterior ROIs to conjunction-based in more anterior ROIs, rather than effects localized to specific, neatly circumscribed regions. In all analyses, data from left and right hemispheres were combined into a single ROI. A subset of representative ROIs was selected for analysis, namely, V1, V2, V3, LOC, PRC, PHC, and HC. LOC was chosen as an intermediate station primarily for its well-established role in representing local contours and object form, without having been routinely implicated in recognition memory (Malach et al. 1995; Grill-Spector et al. 2001; Güçlü and van Gerven 2015).

#### fMRI Memory Score Analysis

To analyze test phase fMRI data, for each separate test run we constructed a generalized linear model with ten regressors: correct responses to each of Familiar, Recombination and Novel stimulus types; incorrect responses; and six motion nuisance regressors. Each non-motion regressor combined a variable boxcar model of the stimulus time-series, with the boxcar beginning at stimulus onset and ending at response time, convolved with a canonical hemodynamic response function. For each participant, the model estimated activation as beta weights (*β*) for every voxel in each of the four conditions.

Memory signal indices were derived from two contrasts, defined in (1) and (2). To index feature memory, we contrasted beta weights from Novel and Recombination correct trials because the novelty/familiarity of individual features is the mnemonic factor discriminating these trial types (both Novel and Recombination stimuli possess a novel three-feature conjunction, but only Novel stimuli possess novel features). To index conjunction memory, we contrasted the beta weights from Recombination and Familiar correct trials because the novelty/familiarity of the conjunction is the mnemonic factor discriminating these trial types (both Recombination and Familiar stimuli possess familiar features, but only Recombination stimuli possess a novel three-feature conjunction). For feature memory, the beta weights for correct Recombination trials (*β_Recombination_*) were subtracted from the beta weights for correct Novel trials (*β_Novel_*). For conjunction memory, the beta weights for correct Familiar trials (*β_Familiar_*) were subtracted from the beta weights for correct Recombination trials (*β_Recombination_*).

For both memory contrasts we obtained a directional Cohen’s *d* effect size, to allow comparison of the magnitude of effects across ROIs with different numbers of voxels and different signal-to-noise levels (Rosenthal 1994; Lakens 2013; Erez et al. 2016). Note that betaweights were calculated *per run* in each voxel individually. Thus, to obtain Cohen’s *d* for a given ROI, betaweights were first averaged across all voxels within the ROI, and then Cohen’s *d* was taken as the mean difference in average betaweights between the two conditions (paired by run), divided by the standard deviation of this difference across runs. Hence:

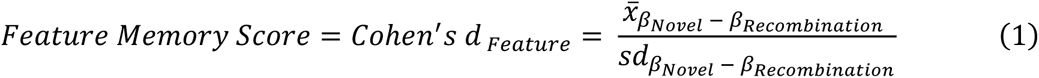

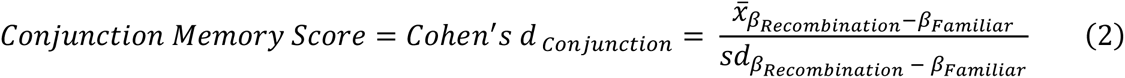

Where *β_Novel_* − *β_Recombination_* is a subscript denoting the difference of two vectors: *β_Recombination_* is a vector of beta weights corresponding to “Novel” trials that were responded to correctly, from all runs; and *β_Recombination_* is a vector of beta weights for correct “Recombination” trials, from all runs. Thus, 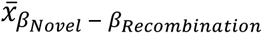 is the mean *β_Novel_* − *β_Recombination_* difference vector for an ROI, and 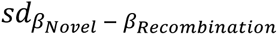 is the standard deviation of the *β_Novel_* − *β_Recombination_* difference vector for that ROI. Analogously, *β_Recombination_* − *β_Familiar_* denotes the difference of two vectors: *β_Recombination_* is a vector from all runs for correct “Recombination” trials; and *β_Familiar_* is a vector from all runs for correct “Familiar” trials. Thus, 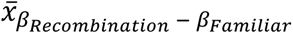 is the mean *β_Recombination_* − *β_Familiar_* difference vector for an ROI, and 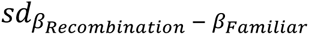 is the standard deviation of the *β_Recombination_* − *β_Familiar_* difference vector.

#### Correlation of Neural Memory Scores and Behavioral Memory Performance

We assessed recognition memory performance using *d’*. Because there were three trial types (and three possible responses), Familiar, Recombination and Novel, *d’* was calculated for each possible pairwise comparison of trial types, yielding four *d’* scores (Table 1). In each pairwise comparison, trials of the third outstanding trial type were removed from the analysis. Any (erroneous) responses to the two remaining trial types that invoked the outstanding trial type were binned as a hit, miss, false alarm or correct rejection, depending on the comparison.

**Table 1:** Discriminability of Mnemonic Stimulus Classes for Fribbles.

| Comparison Condition | $d'$ | Trial Types | Hit/Correct Rejection | Miss/False Alarm |
| --- | --- | --- | --- | --- |
| Familiar vs. Novel (I) | 4.44 | Familiar | "Familiar" & "Recombination" | "Novel" |
|  |  | Novel | "Novel" | "Familiar" & "Recombination" |
| Familiar vs. Novel (II) | 3.22 | Familiar | "Familiar" | "Novel" & "Recombination" |
|  |  | Novel | "Novel" & "Recombination" | "Familiar" |
| Familiar vs. Recombination<br>*a.k.a. Conjunction Memory $d'$* | 2.26 | Familiar | "Familiar" | "Recombination" & "Novel" |
|  |  | Recombination | "Recombination" & "Novel" | "Familiar" |
| Recombination vs. Novel<br>*a.k.a. Feature Memory $d'$* | 4.33 | Recombination | "Recombination" & "Familiar" | "Novel" |
|  |  | Novel | "Novel" | "Recombination" & "Familiar" |

Two of the four *d’* scores were analogous to the neural feature and conjunction memory scores – Novel vs. Recombination and Recombination vs. Familiar. For these comparisons, we measured the correlation between neural memory scores and recognition memory *d’* scores. For feature memory, we used the Recombination vs. Novel comparison: only Recombination and Novel trials were included, and a “Familiar” response counted as a hit for Recombination stimuli, but a false alarm for Novel stimuli. For conjunction memory, we used the Familiar vs. Recombination comparison: only Familiar and Recombination trials were included, and a “Novel” response counted as a correct rejection for Recombination stimuli, but a miss for Familiar stimuli.

#### Bayes Factor Analyses

For analyses in which the data allowed for *parametric* statistical tests, we replaced traditional frequentist tests with Bayes Factors (BFs), to assess statistical reliability. Bayesian hypothesis testing offers several advantages over null hypothesis significance testing, including that BFs quantify the relative evidence of the alternative hypothesis over the null (rather than simply specifying the null and determining whether to reject it); they allow quantitative assessment of the likelihood that the null hypothesis is true (rather than just determining whether it can be rejected); they are not biased against the null hypothesis; and interpretation of a BF is unaffected by the sampling intentions of the experimenter (Wagenmakers et al. 2018).

A BF is the relative probability of observed data (*D*) under two alternative theoretical claims (e.g., *M_1_, M_2_*). Although any two hypotheses can be compared, here it is a likelihood ratio of the data under the assumption of the presence of an effect or the absence of an effect (BF = P [D | M1] / P [D | M2]; see Schmalz et al. 2021 for a detailed overview). For example, when testing the significance of a positive correlation, the BF compares the likelihood of the observed data under the alternative hypothesis that the true linear correlation is greater than 0, to the likelihood of the observed data under the null hypothesis that the true linear correlation is equal to or less than 0. In general, BFs close to 1 indicate that the data are equally likely under both hypotheses, while values much larger than 1 or much smaller than 1 indicate evidence favoring one hypothesis over the other. Categorical guidelines for the interpretation of BFs suggest that values greater than 3 indicate “moderate” or “substantial” evidence for a hypothesis, greater than 10 “strong evidence”, greater than 30 “very strong” evidence, and greater than 100 “decisive” or “extreme” evidence (the same guidelines can be applied to the hypothesis in the denominator by making these values fractions; Jeffreys 1961; Raftery 1995).

We used the *BayesFactor* R package (Morey et al. 2015) with default input values for the number of Monte Carlo simulations (10,000) and for the medium scale values of priors (ANOVA: ^1^/_2_ g-prior; *t*-test: 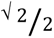 Cauchy prior; correlation: ^1^/_3_ beta prior). The null hypothesis lies in the denominator of the BF, therefore values larger than 1 provide evidence for the alternative hypothesis, e.g., for correlation *t*-tests, BF>1 provides evidence for the alternative hypothesis that there is a significant positive relationship between the two variables. Where we report a BF in the form “^1^/_x_”, x is the BF in standard format (alternative model in the numerator, null in the denominator) but x is <1, indicating greater evidence for the null, so we invert the BF to report evidence for the null. For ANOVA designs with nested models (i.e., a full model containing all main effects and their interactions, as well as restricted models with fewer effects) all models are compared against the null (i.e., a model with only random subject effects). Because BFs from these nested model comparisons share a common denominator, we can directly compare nested models by taking the ratio of their BFs.

#### Nonparametric Permutation Tests

For analyses in which nonparametric tests were required by the structure of the data we used permutation and shuffling to generate nonparametric null distributions and subsequently assessed whether the statistic produced by the true (unshuffled) data fell in the extreme (>.95) of the generated distribution.

We used such nonparametric tests to ask whether the linear trend in memory signal (Cohen’s *d*) across the ventral visual-MTL pathway differed between feature and conjunction memory. In this analysis, we took the feature and conjunction Cohen’s *d* values within each ROI (V1, V2, V3, LOC, PRC, PHC, HC), multiplied these by weights describing opposite linear trends for feature memory (decreasing from early visual areas to MTL) and conjunction memory (increasing from early visual areas to MTL) and summed the resulting products. Next, we generated null distributions for each ROI by shuffling the ROI labels associated with each participant’s set of seven Cohen’s *d* values and repeating this 10,000 times. Using the resulting seven ROI null distributions, we performed the difference of linear trends computation described above 10,000 times, to generate a null distribution for the difference of linear trends. If the true trends in correlation across ROIs are similar for feature and conjunction memory, the Cohen’s *d* values multiplied by opposite-trending weights should sum to zero, but if there are opposite trends in correlation across ROIs, the sum should be greater than zero. We conducted further permutation analyses to test whether each linear trend in isolation was significant, i.e., to ask separately whether there is a significant descending linear trend for feature memory and ascending linear trend for conjunction memory.

Similarly, we evaluated the pattern of correlations across ROIs by testing whether the linear trends in correlation coefficients significantly differed between feature and conjunction memory. For this, we took the inverse normal transformation of correlations within each ROI, multiplied these by weights describing opposite linear trends for feature memory (i.e., decreasing from early visual areas to MTL) and conjunction memory (i.e., increasing from early visual areas to MTL), and summed the resulting products. Next, we generated null distributions for each ROI by shuffling participant Cohen’s *d* and *d’* values, computing their correlation, taking the inverse normal transformation of the correlation score, and repeating this 10,000 times. Using the resulting seven ROI null distributions, we performed the difference of linear trends computation 10,000 times, to generate a null distribution for the difference of linear trends. If the true trends in correlation across ROIs are similar for feature and conjunction memory, the transformed correlation values multiplied by opposite-trending weights should sum to zero; however, if there are opposite trends, the sum should be greater than zero. Finally, we tested each linear trend in isolation: descending for feature memory and ascending for conjunction memory.

## Results

### Behavioral Analysis

Performance on the 1-back task during the study phase, measured as the discriminability between same and different trials (*d’*), was good, with a mean *d’* of 3.21 for Fribbles and 3.26 for Scenes.

Memory discrimination performance in the test phase (Tables 1 and 2) was also good for all comparison types (Familiar vs. Novel (I); Familiar vs. Novel (II); Familiar vs. Recombination; Recombination vs. Novel). For Fribbles and Scenes separately, we explored differences in memory discrimination performance with a BF analysis akin to a one-way ANOVA. For both stimulus sets, we found extreme evidence for a main effect of comparison type, indicating that *d’* differed between pairwise comparisons (BF > 150). As expected, *d’* scores were lowest, but still above chance, in the comparison of Familiar and Recombination trials – trials in which all features were familiar, making mnemonic discrimination difficult.

**Table 2:**
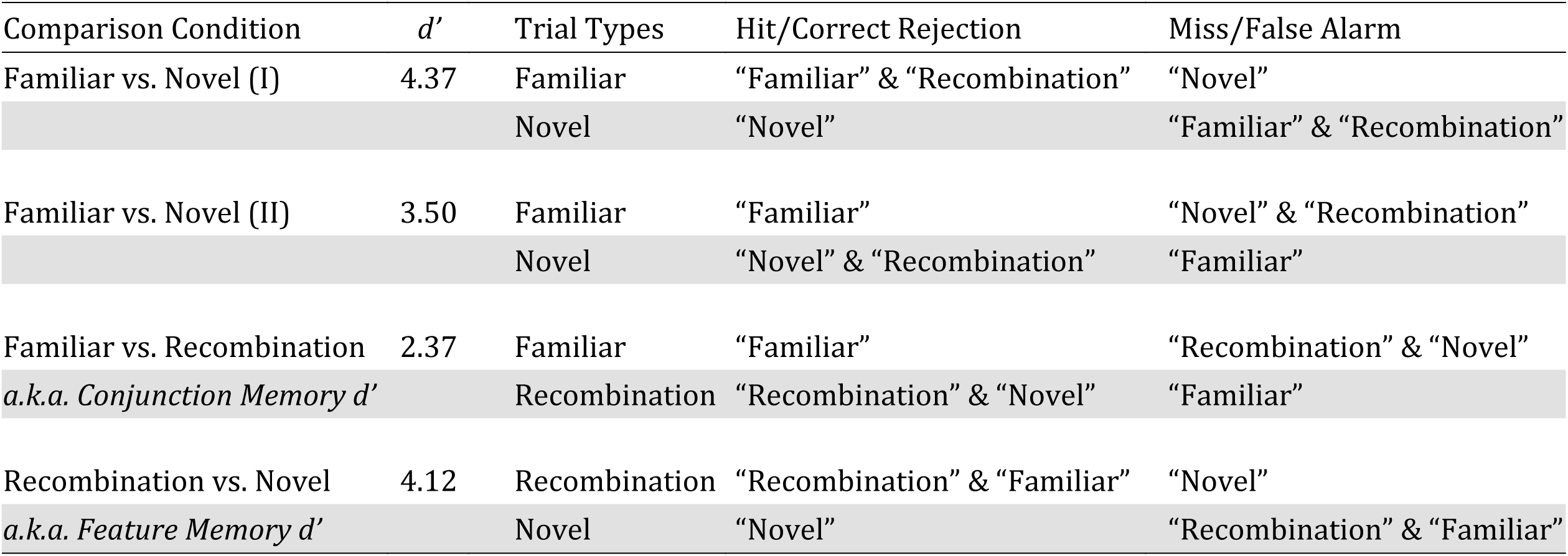
Discriminability of Mnemonic Stimulus Classes for Scenes.

### fMRI Memory Score Analysis

To interpret the neural memory scores (Cohen’s *d*), for each stimulus set, we computed BFs to compare nested models in an analysis equivalent to a two-way (7×2) ANOVA with factors of ROI (V1, V2, V3, LOC, PRC, PHC, HC) and memory score type (Feature Memory, Conjunction Memory). Memory scores are shown in Figures 5 and 6 (see Supplementary Figures S2 through S5 for root mean beta weights and individual data points).

**Figure 5.**
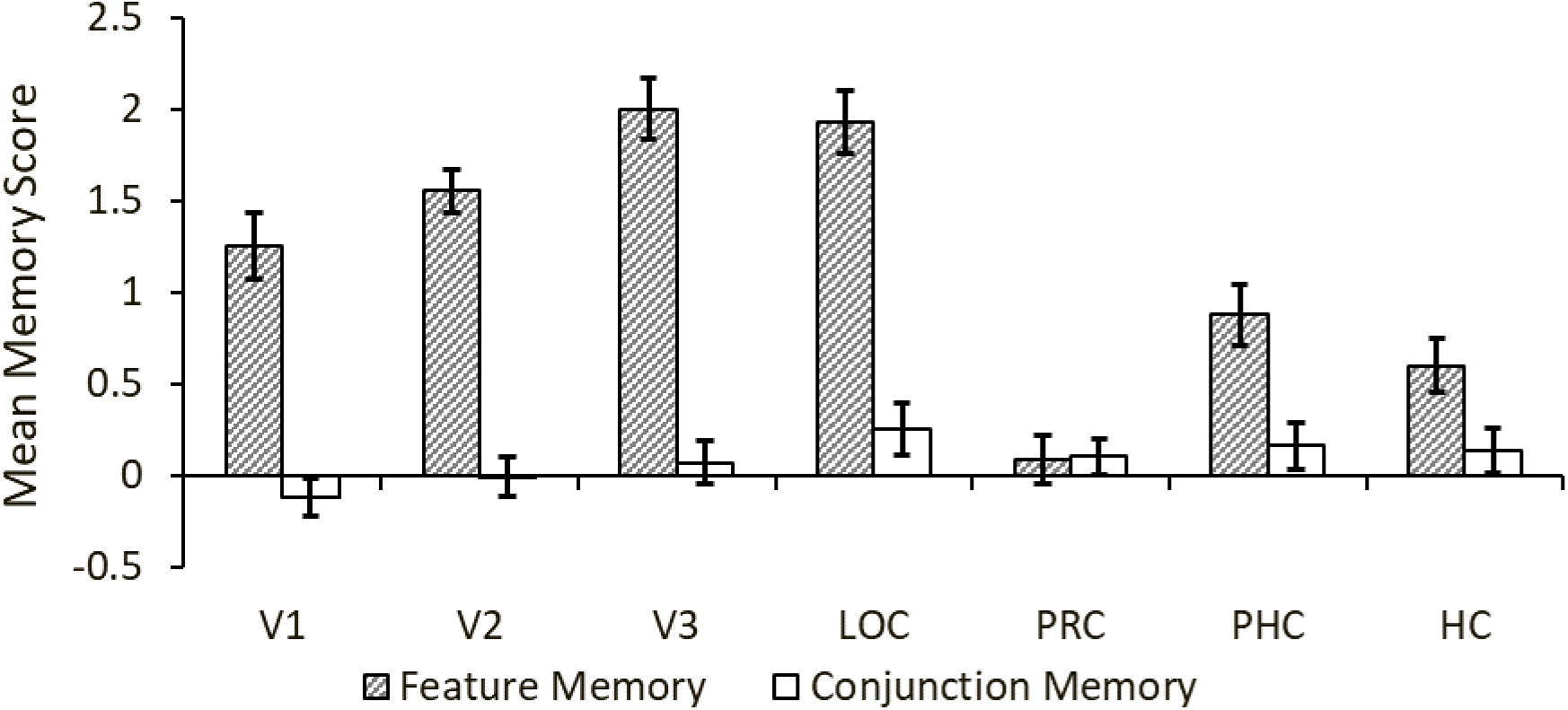
Memory Scores for Fribbles. Memory scores were derived by calculating the effect size, measured by Cohen’s *d*, for the contrast of Novel versus Recombination trials (Feature Memory) and for the contrast of Recombination versus Familiar trials (Conjunction Memory). Error bars are within-subject SEM.

**Figure 6.**
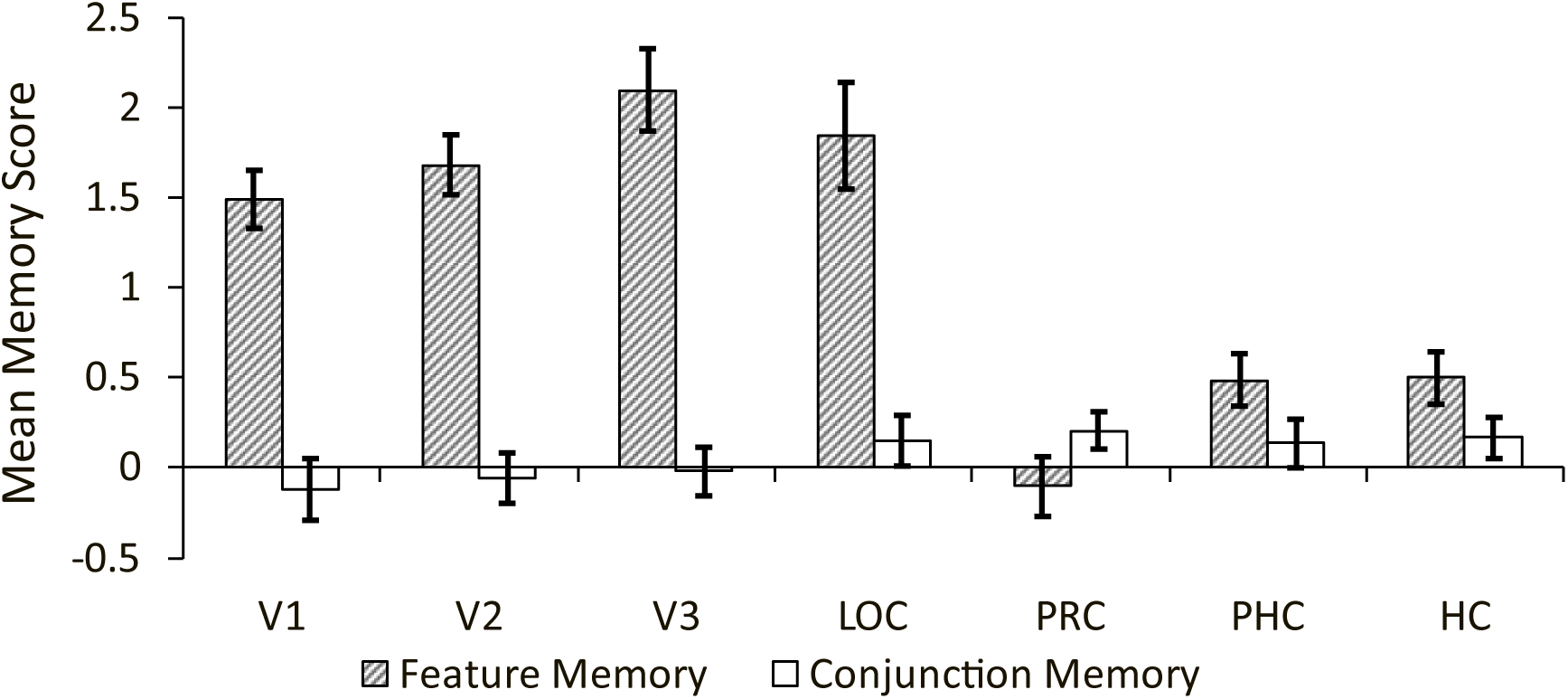
Memory Scores for Scenes. Memory scores were derived by calculating the effect size, measured by Cohen’s *d*, for the contrast of Novel versus Recombination trials (Feature Memory) and for the contrast of Recombination versus Familiar trials (Conjunction Memory). Error bars are within-subject SEM.

For Fribbles, BFs reflected extreme evidence in favor of the interactive model (ROI x memory type) over the null (BF > 150) and over the next two most favored models (additive [ROI + memory type]: BF = 1.16 x 10^43^/ 3.98 x 10^32^ > 150; memory type only: BF = 1.16 x 10^43^/ 4.41 x 10^24^ > 150). Numerically, the signal for feature memory was greatest in ROIs in early and mid-ventral visual stream (V1, V2, V3 and LOC) and decreased moving anteriorly toward MTL, whereas the conjunction memory signal was greatest in late-stage ventral visual stream (LOC) and MTL (PHC, PRC, HC) and decreased moving posteriorly. We explored the interaction by separating the data for feature and conjunction memory and performing two separate model comparisons, finding extreme evidence for a main effect of ROI for feature memory (BF > 150) and moderate evidence against a main effect of ROI for conjunction memory (BF = 1/.27 = 3.71; but see the linear trend analysis, below).

Conjunction memory scores for Fribbles clustered near zero, and so we tested whether any significantly exceeded zero, in line with our *a priori* predictions. For Fribbles, there were five ROIs with numerically positive conjunction memory scores, all relatively anterior (V3, LOC, PRC, PHC, HC). The alternative hypothesis, H_1_, was defined as Cohen’s D > 0, and the null hypothesis, H_0_, as Cohen’s D <= 0. We found either anecdotal evidence in favor of the null (i.e., conjunction memory <= 0; BF _V3_ = 1/.34 = 2.95; BF _PRC_ = 1/.45 = 2.23) or almost no evidence for either the alternative or the null (BF _LOC_ = 1/.96 = 1.04; BF _PHC_ = 1/.70 = 1.42; BF _HC_ = 1/.61 = 1.63). It is instructive that in all anterior ROIs (i.e., LOC, PRC, PHC and HC), conjunction memory was numerically greater than zero, but we found neither substantial evidence for, nor substantial evidence against, the presence of conjunction memory. Because BFs can impart evidence for both the alternative and the null, the lack of evidence either way suggests insufficient power to detect conjunction memory signals at the group level with Fribbles. This was not the case with Scenes (see below).

We next used nonparametric permutation tests to ask whether there were differing linear trends in the memory signals across the ventral visual-MTL (descending for features and ascending for conjunctions), as described in Materials and Methods. For Fribbles, we included weights for just six ROIs, omitting HC, and we weighted PRC and PHC equally (feature memory [3 2 1 -1 -2.5 -2.5]; conjunction memory [-3 -2 -1 1 2.5 2.5]). We excluded HC for the Fribbles dataset because we did not predict, a priori, that object conjunctions would be represented as far anterior as HC, and indeed we had no predictions for the type of code (feature versus conjunction) that HC might contribute for objects. We weighted PRC and PHC equally because both form part of the parahippocampal gyrus and project to HC via the entorhinal cortex (Naber et al. 1997; Burwell 2000). The true weighted linear sum was significantly greater than zero for Fribbles data (*p* < .001), indicating that the linear trend of Cohen’s *d* across ROIs differed for feature and conjunction memory. We conducted further permutation analyses to test whether each linear trend in isolation (descending for feature memory; ascending for conjunction memory) was significant. For Fribbles, the descending linear trend for feature memory was highly significant (p < .001) and the ascending linear trend for conjunction memory was borderline significant (p = .0498).

For Scenes, BFs reflected extreme evidence in favor of the interactive model (ROI x memory type) over the null (BF > 150) and over the next two most favored models (additive [ROI + memory type]: BF = 1.74 x 10^35^/ 4.91 x 10^22^ > 150; memory type only: BF = 1.74 x 10^35^/ 2.37 x 10^17^ > 150). Numerically, the signal for feature memory was greatest in ROIs in early and mid-ventral visual stream (V1, V2, V3 and LOC) and decreased moving anteriorly toward MTL (Figure 6); the greatest signal for conjunction memory occurred in MTL (PHC, PRC, HC) and late-stage ventral visual stream (LOC) and decreased moving posteriorly toward early visual areas. We explored this interaction by separating the data for feature and conjunction memory and performing two separate model comparisons examining the effect of ROI. We found extreme evidence for a main effect of ROI for feature memory (BF > 150) and anecdotal evidence against a main effect of ROI for conjunction memory (BF = 1/.47 = 2.13; but see the linear trend analysis, below).

Conjunction memory scores for Scenes also clustered near zero, and thus we tested whether they exceeded zero in ROIs with positive scores (again, H_1_ was Cohen’s D > 0 and H_0_ was Cohen’s D <=0). Of the four ROIs with numerically positive conjunction memory scores (LOC, PRC, PHC and HC), there was anecdotal evidence in favor of the null in LOC (i.e., conjunction memory <= 0; BF _LOC_ = 1/.50 = 2.02), a lack of evidence for either the alternative or the null in PHC and HC (BF _PHC_ = 1/.55 = 1.82; BF _HC_ = 1.27), and substantial evidence in favor of conjunction memory scores greater than zero in PRC (BF _PRC_ = 5.68).

In a final assessment of Cohen’s *d* for Scenes, we used nonparametric permutation tests to ask whether feature and conjunction memory signals exhibited different linear trends across the ventral visual-MTL pathway (descending for features and ascending for conjunctions) as described in Materials and Methods. For Scenes, we included weights for all seven ROIs (i.e., including HC), with PRC and PHC weighted equally, resulting in feature weights of [3 2 1 0 -1.5 -1.5 -3] and conjunction weights of [-3 -2 -1 0 1.5 1.5 3]. The true weighted linear sum of the difference in linear trends was significantly greater than zero for Scenes (*p* < .001), indicating that the linear trends across ROIs differed for feature and conjunction memory signals. Examining each linear trend in isolation, both the descending linear trend for feature memory (p < .001) and the ascending linear trend for conjunction memory (p = .02) were significantly different from zero.

### Correlation of Neural Memory Scores and Behavioral Memory Performance

To assess whether the observed neural signals for feature and conjunction memory in Scenes are implicated in recognition memory decisions, we correlated neural memory scores (Cohen’s *d*) with recognition memory performance (*d*’) across participants, in each ROI. Specifically, we correlated (1) conjunction memory scores with conjunction memory *d’* (both obtained by comparing Familiar and Recombination trials) and (2) feature memory scores with feature memory *d’* (both obtained by comparing Novel and Recombination trials) for each ROI (V1, V2, V3, LOC, PRC, PHC, HC), separately.

In Fribbles, Pearson correlation coefficients were numerically as predicted by a representational account: greatest in posterior areas of the ventral visual-MTL pathway for feature memory, and greatest in more anterior regions, LOC and PRC, for conjunction memory (Figure 7). We tested whether the linear trends in correlation coefficients for Fribbles significantly differed between feature and conjunction memory. As with the linear trends in Cohen’s *d*, for Fribbles we omitted HC and set weights for PRC and PHC equal (feature memory: [3 2 1 -1 -2.5 -2.5]; conjunction memory: [-3, -2 -1 1 2.5 2.5]). The true weighted linear sum of the difference in trends was borderline significantly greater than zero (p = .06). We also examined the linear trend for each memory type in isolation in Fribbles: there was no significant linear trend for either feature memory (*p* = .10) or conjunction memory (*p* = .18). Together, these results – which are numerically in line with predictions and with the results for Scenes (see below) – accord with the notion that the experimental design for Fribbles stimuli was underpowered.

**Figure 7.**
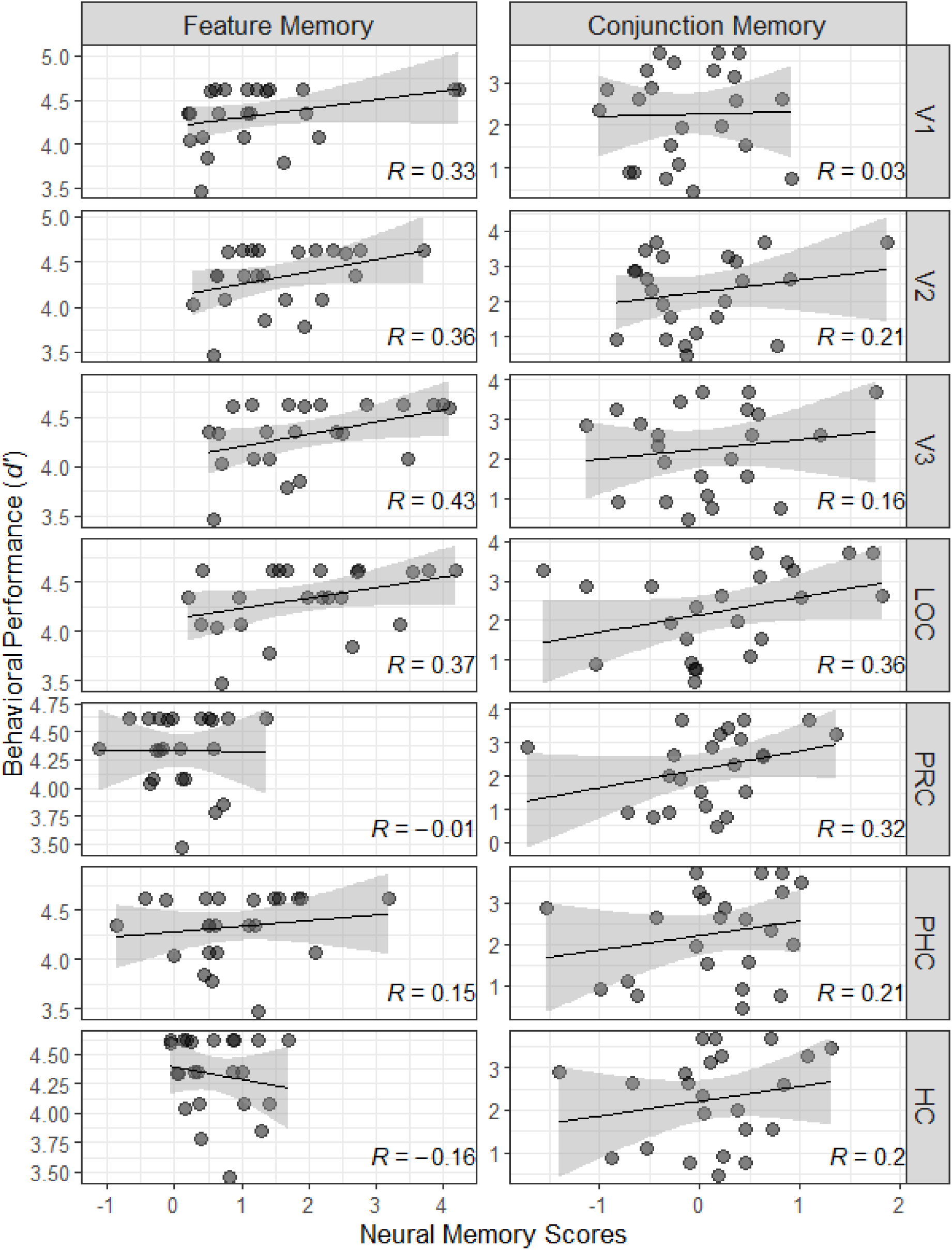
Correlations between Memory Scores and *d’* for Fribbles. Memory scores derived from effect sizes (Cohen’s *d*) in the neural data were correlated with *d’* scores from corresponding behavioral data for Feature Memory (*left* column; Novel versus Recombination trials) and Conjunction Memory (*right* column; Recombination versus Familiar trials). Regression lines are displayed with 95% confidence intervals. Pearson’s R is shown in each panel. HC was not included in linear trend analyses for Fribbles (see text) but is shown here for completeness.

Finally, in Fribbles, we computed BFs to assess the evidence for a positive brain-behavior correlation in each ROI. For each ROI, H_1_ was defined as Pearson’s R > 0, and H_0_ as Pearson’s R <= 0. Feature memory correlations yielded the largest BFs in the four posterior, visual brain regions, where evidence was strong (Table 3). In contrast, the MTL ROIs showed either anecdotal evidence in favor of a positive feature memory correlation (PHC), no evidence in favor or against a positive correlation (PRC), or anecdotal evidence in favor of the null (HC). For conjunction memory, the evidence for a brain-behavior correlation in Fribbles was strong in LOC (13.36), an intermediate visual region known for processing objects holistically (e.g., Malach et al. 1995; Grill-Spector et al. 1998), and approaching strong in PRC (9.73), another region heavily implicated in whole-object processing (e.g., Buckley et al. 2001; Barense et al. 2007).

**Table 3:**
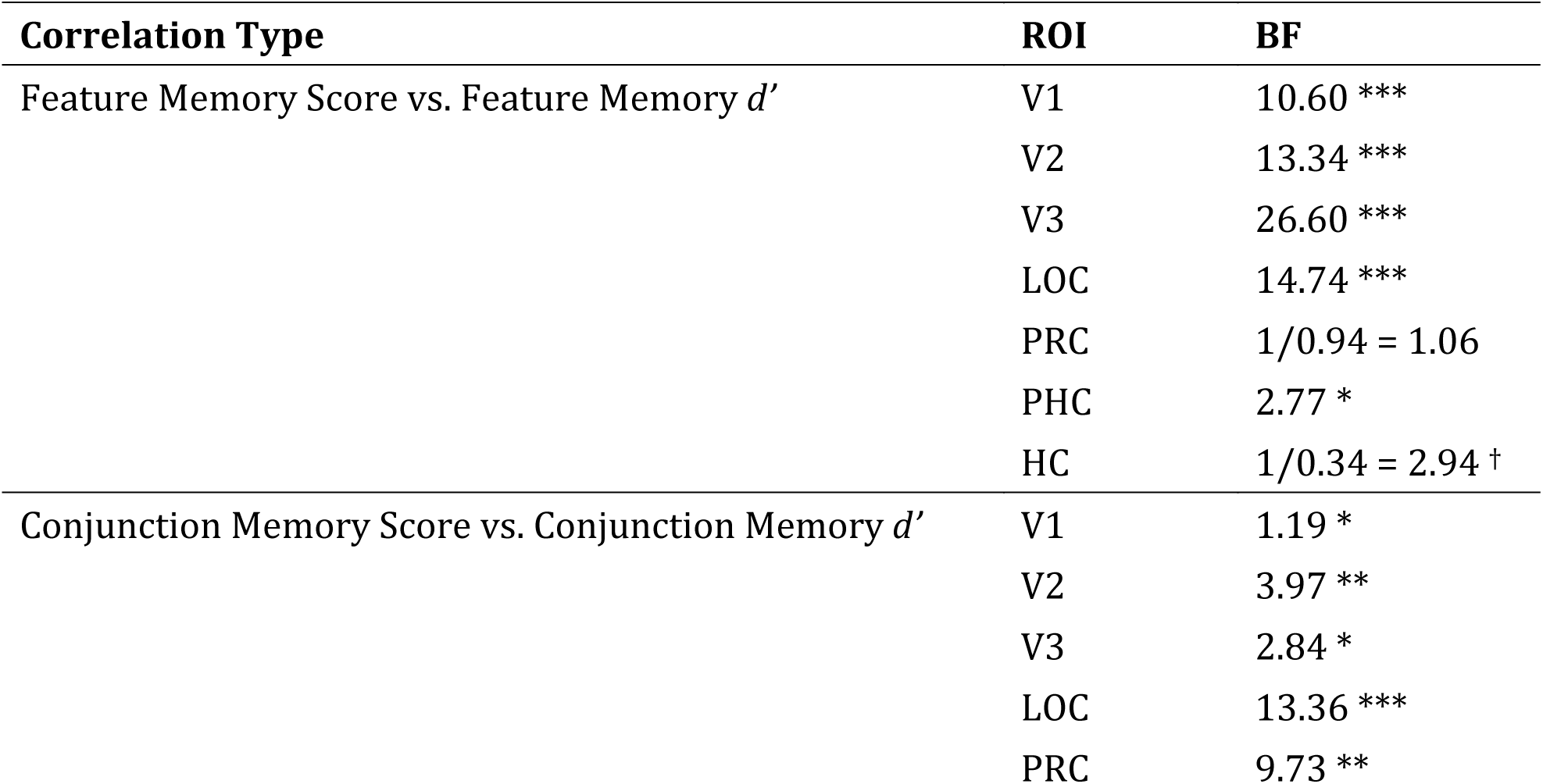
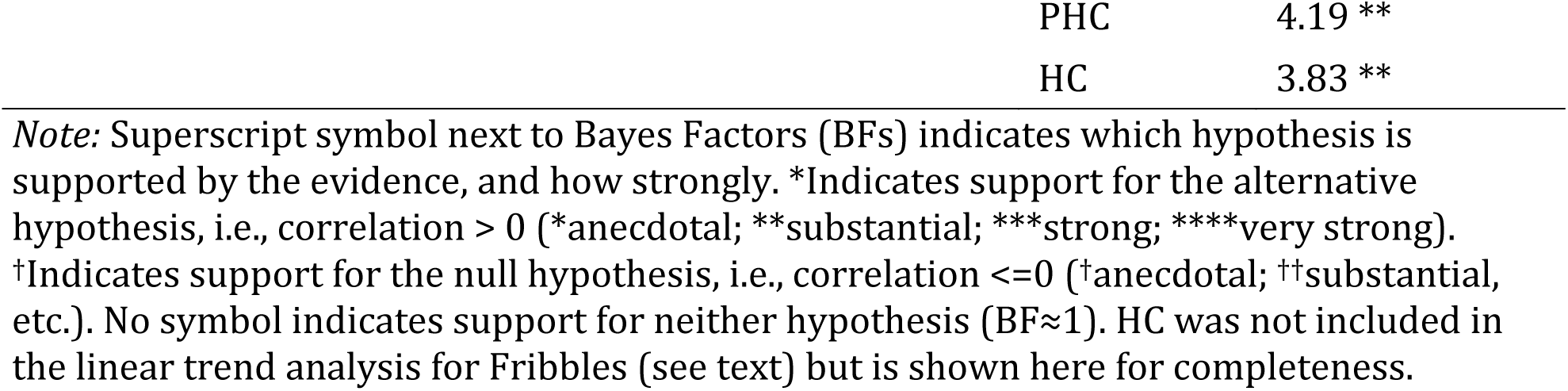
Bayes Factors for Correlations between Memory Scores and *d’* for Fribbles.

| Correlation Type | ROI | BF |
| --- | --- | --- |
| Feature Memory Score vs. Feature Memory $d'$ | V1 | 10.60 *** |
|  | V2 | 13.34 *** |
|  | V3 | 26.60 *** |
|  | LOC | 14.74 *** |
|  | PRC | 1/0.94 = 1.06 |
|  | PHC | 2.77 * |
|  | HC | 1/0.34 = 2.94 † |
| Conjunction Memory Score vs. Conjunction Memory $d'$ | V1 | 1.19 * |
|  | V2 | 3.97 ** |
|  | V3 | 2.84 * |
|  | LOC | 13.36 *** |
|  | PRC | 9.73 ** |

|  |  |
| --- | --- |
| PHC | 4.19 ** |
| HC | 3.83 ** |

*Note:* Superscript symbol next to Bayes Factors (BFs) indicates which hypothesis is supported by the evidence, and how strongly. \*Indicates support for the alternative hypothesis, i.e., correlation > 0 (\*anecdotal; \*\*substantial; \*\*\*strong; \*\*\*\*very strong). †Indicates support for the null hypothesis, i.e., correlation ≤ 0 (†anecdotal; ††substantial, etc.). No symbol indicates support for neither hypothesis (BF≈1). HC was not included in the linear trend analysis for Fribbles (see text) but is shown here for completeness.

For Scenes, Pearson correlation coefficients for feature memory were greatest in posterior areas of the ventral visual-MTL pathway, and correlation coefficients for conjunction memory were greatest in MTL regions (Figure 8). We evaluated the pattern of correlations across ROIs by testing whether the linear trends in correlation coefficients significantly differed between feature and conjunction memory (see Materials and Methods). As with the linear trends in Cohen’s *d* for Scenes, we included weights for all seven ROIs with PRC and PHC weighted equally (feature memory: [3 2 1 0 -1.5 -1.5 -3]; conjunction memory: [-3 -2 -1 0 1.5 1.5 3]). The true weighted linear sum of the difference in linear trends was significantly greater than zero for Scenes (*p* = .007), indicating that the linear trend of correlation coefficients across ROIs differed for feature and conjunction memory. We next examined the linear trend in each memory type in isolation, finding a borderline significant linear trend in the descending direction for feature memory (*p* = .05) and a significant linear trend in the ascending direction for conjunction memory (*p* = .03).

**Figure 8.**
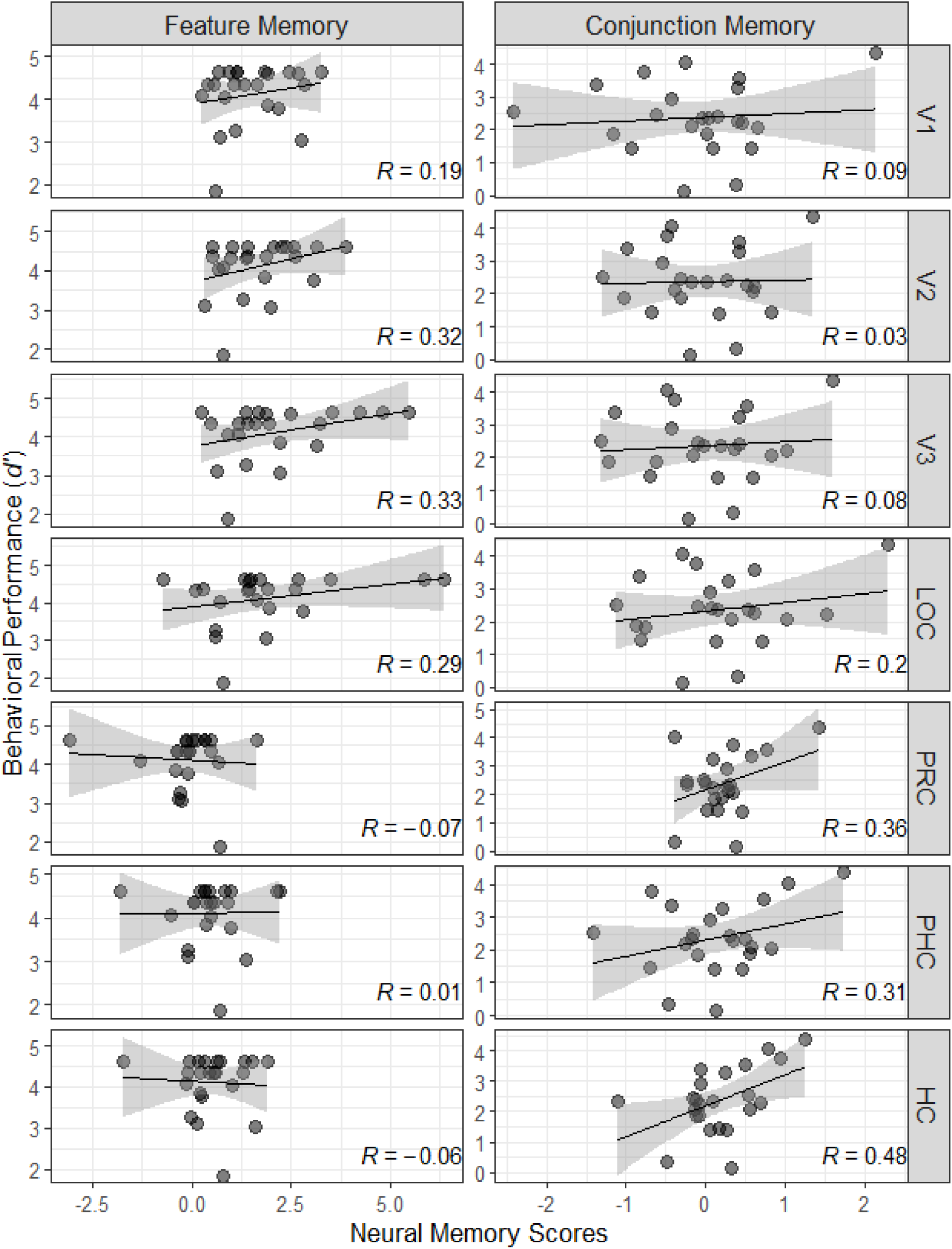
Correlations between Memory Scores and *d’* for Scenes. Memory scores (Cohen’s *d*) in the neural data were correlated with *d’* scores from corresponding behavioral data for Feature Memory (*left* column; Novel versus Recombination trials) and Conjunction Memory (*right* column; Recombination versus Familiar trials). Regression lines are displayed with 95% confidence intervals. Pearson’s R is shown in each panel.

In a final assessment of the relationship between neural memory scores and behavioral memory performance for Scenes, we calculated BFs to assess the evidence for a positive correlation in each ROI for each memory type (Table 4). Again, H_1_ was defined as Pearson’s R > 0, and H_0_ as Pearson’s R <= 0. For feature memory, the strongest evidence for a brain-behavior correlation was found in posterior, visual ROIs, especially V2 and V3. In MTL regions, there was a lack of evidence to weak anecdotal evidence in favor of the null, i.e., correlation <= 0. For conjunction memory, evidence for a correlation was absent to anecdotal in posterior regions but increased along the pathway until it reached “very strong” in HC, suggesting that the conjunction memory signals in HC are the ones most related to the mnemonic discrimination of scene conjunctions.

**Table 4:**
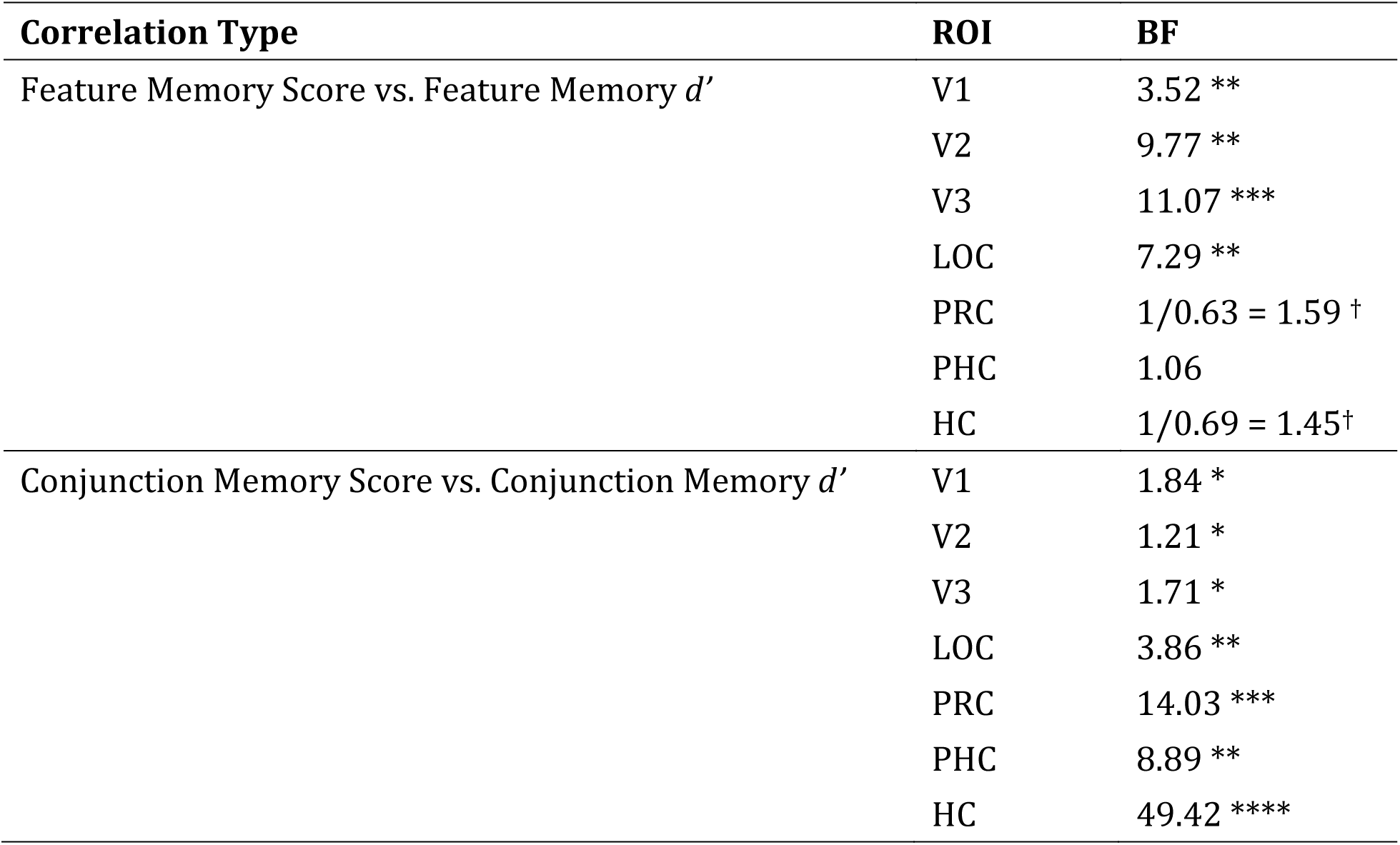
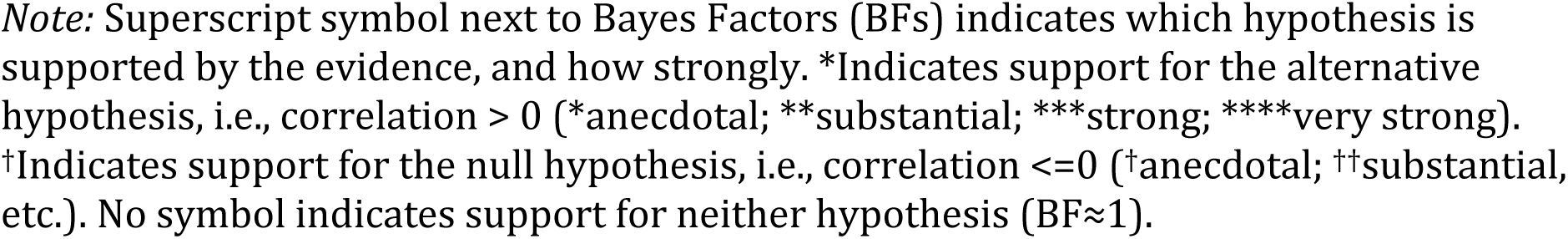
Bayes Factors for Correlations between Memory Scores and *d’* for Scenes.

| Correlation Type | ROI | BF |
| --- | --- | --- |
| Feature Memory Score vs. Feature Memory $d'$ | V1 | 3.52 ** |
|  | V2 | 9.77 ** |
|  | V3 | 11.07 *** |
|  | LOC | 7.29 ** |
|  | PRC | 1/0.63 = 1.59 <sup>†</sup> |
|  | PHC | 1.06 |
|  | HC | 1/0.69 = 1.45<sup>†</sup> |
| Conjunction Memory Score vs. Conjunction Memory $d'$ | V1 | 1.84 * |
|  | V2 | 1.21 * |
|  | V3 | 1.71 * |
|  | LOC | 3.86 ** |
|  | PRC | 14.03 *** |
|  | PHC | 8.89 ** |
|  | HC | 49.42 **** |

*Note:* Superscript symbol next to Bayes Factors (BFs) indicates which hypothesis is supported by the evidence, and how strongly. \*Indicates support for the alternative hypothesis, i.e., correlation > 0 (\*anecdotal; \*\*substantial; \*\*\*strong; \*\*\*\*very strong). †Indicates support for the null hypothesis, i.e., correlation ≤ 0 (†anecdotal; ††substantial, etc.). No symbol indicates support for neither hypothesis (BF≈1).

## Discussion

We tested the hypothesis that the neural signals supporting recognition memory shift within the brain according to the complexity of information that participants are asked to recognize. This hypothesis stems from representational accounts, in which the neuroanatomical organization of perception and memory is governed by the information each brain region represents, rather than – as under a “multiple memory systems” view – by the perceptual or mnemonic function each region performs. In this neuroimaging study, we used a recognition memory task that assessed mnemonic discrimination at two levels, one requiring simpler, feature-based representations and the other more complex, conjunction-based representations. These levels were created by testing participants’ memory for visual stimuli constructed from conjunctions of pre-defined, binary features. We assayed simple memory by contrasting stimuli that were discriminable only by the familiarity of their features, and we measured more complex memory by contrasting stimuli that were discriminable only by the familiarity of their conjunctions.

We report three main findings. First, the strongest neural signals for long-term (i.e., longer than 30 seconds) mnemonic discrimination of visual features lie outside of MTL, in visual cortex. Second, the locus of neural recognition memory signals shifted with the complexity of the stimulus representations required for the mnemonic discrimination: the strongest feature memory signals lay posterior to the strongest conjunction memory signals (when they were detected, i.e., for scenes) within the ventral visual-MTL pathway. Third, these memory signals predicted participants’ memory performance, and the strongest brain-behavior correlations for feature-based memory emerged in posterior, visual regions, whereas the strongest correlations for conjunction-based memory emerged in anterior regions, mostly MTL. Linear trend analyses of these brain-behavior correlations revealed trends of opposite direction for feature versus conjunction memory, in the Scenes dataset (and a numerically similar pattern in Fribbles that was borderline significant).

The locations of the neural memory signals and the correlations of those signals with mnemonic performance indicate that the ventral visual-MTL pathway supports recognition memory behavior as the Representational-Hierarchical account predicts it should. Moreover, as predicted by representational accounts, the cortical sites most associated with feature-based versus conjunction-based *memory* performance are the same sites previously found to support feature-based versus conjunction-based visual *perception*. We highlight here three findings that illustrate this claim. First, we observed in both objects (Fribbles) and Scenes that evidence for a correlation between feature-based memory signals and feature-based mnemonic discrimination was substantial or strong in visual regions (V1 to LOC), and anecdotal or absent in MTL regions, which accords with findings from perceptual studies that feature-based representations reside in early visual regions (e.g., Hubel and Wiesel 1965; Kobatake and Tanaka 1994; Mazer et al. 2002; Kamitani and Tong 2005; Henriksson et al. 2008; Brouwer and Heeger 2009; Serences, Saproo, et al. 2009). Second, for Fribbles – which are novel, visual objects without semantic meaning, but defined by their unique conjunction – we found the strongest evidence for a brain-behavior correlation for conjunction memory in LOC (BF=13.36) and PRC (BF=9.73). LOC and PRC are the two regions among those we tested that have been most clearly implicated in holistic visual object-processing as measured with perceptual tasks (e.g., Malach et al. 1995; Grill-Spector et al. 1998; Barense et al. 2007). Third, for Scenes, evidence for a conjunction memory correlation was absent to anecdotal in early visual cortex (V1, V2, V3) and increased with anterior progression, reaching a maximum later in the pathway than for Fribbles, in HC. These patterns accord with previous findings that complex conjunctive representations reside in anterior sites during perceptual tasks (e.g., Desimone et al. 1984; Kobatake and Tanaka 1994; Kanwisher et al. 1997; Kriegeskorte et al. 2008; Ostwald et al. 2008; Drucker and Aguirre 2009; Cowell et al. 2017), and that the perception of scenes engages hippocampus, particularly when the task demands require conjunctive representations (Lee et al. 2005; Lee et al. 2006; Lee et al. 2007; Lee et al. 2008). In sum, the cortical sites most associated with feature- or conjunction-based discrimination in this memory task were the same as those previously found to underpin feature- or conjunction-based processing in perceptual tasks.

To our knowledge, this is the first report of a within-study manipulation of the level of complexity of representations required for a long-term memory discrimination. That is, using the same stimuli and a single task – stimuli comprising binary features combined into conjunctions, along with discrimination between Old, Recombined and Novel memory statuses – we teased apart feature- and conjunction-based memory in terms of both neural signals and behavioral performance. This allowed direct comparison of the two types of memory content, demonstrating that manipulating the level of complexity leads to shifts in the locus of long-term memory signals. Furthermore, the presence of significant brain-behavior correlations for both feature- and conjunction-based memory, in the locations expected based on prior literature, suggests that these BOLD signatures of memory relate to behavioral responses in a long-term recognition memory task.

Many recent empirical studies have demonstrated a critical role for visual cortical representations in visual short-term memory (Harrison and Tong 2009; Serences, Ester, et al. 2009; Christophel et al. 2012; Jerde et al. 2012; Riggall and Postle 2012; Sprague et al. 2014; Christophel et al. 2017). But the length of the retention delay in those studies was on the order of seconds, whereas in the current paradigm, the delay between study and recognition test was a minimum of twenty minutes. This suggests that visual cortex is involved in memory not just over brief periods in which active maintenance is required, but for longer periods in which synaptic changes are likely to underlie the observed behavioral changes (Clarke et al. 2010; Cooke and Bear 2010; Banks et al. 2014; Cooke and Bear 2015).

Another literature that has shown a role for sensory cortices in long-term memory retrieval has couched that role as “cortical reinstatement”, with the reinstatement typically said to be mediated or triggered by the HC (Le Bihan et al. 1993; Eichenbaum 2000; Johnson and Rugg 2007; Bosch et al. 2014; Danker et al. 2017). To be clear, we do not contest the existence of cortical reinstatement, which clearly plays an important role – complementary to the role of signal strength in familiarity judgments – in part-cued retrieval of complex, associative memories. Here, we simply ask whether hippocampally-triggered cortical reinstatement is mandatorily involved in all retrieval, including recognition memory for simple, low-dimensional, visual information. Our interpretation of the present results is guided by parsimony: given that familiarity/novelty signals provide a powerful mechanism for recognition memory (e.g., Rutishauser et al. 2006), and given that in the feature-based memory task part-to-part retrieval of high-dimensional representations was not required, we prefer not to invoke hippocampally triggered reinstatement. In support of this parsimonious interpretation, for feature-based mnemonic discrimination we found that memory signals within MTL regions do not robustly predict behavior, even though signals from these same regions were strongly correlated with behavior for conjunction-based discriminations. That is, in the present data, the involvement of MTL memory signals appears to come and go with manipulations in the content of the memory, just as was found in recent neuroimaging studies of visual recall (Ross et al. 2018; Gardette et al. 2022). Moreover, in Scenes, feature-memory was better predicted by visual cortical signals than by MTL signals (and the same trend was found numerically in Fribbles). This pattern fits the notion that the signals underlying recognition memory lie distributed along the ventral visual-MTL pathway according to the content of the memory, rather than the notion of mandatory hippocampally triggered cortical reinstatement. However, there exist other measures of reinstatement, including multivariate effects or hippocampal-cortical connectivity, which we did not compute. Without investigating these measures, a cortical reinstatement account cannot be ruled out.

We note here three limitations to the present findings. First, the sample size was relatively small (data collection was curtailed by the COVID-19 pandemic), meaning that these results require replication in future work. Second, in Fribbles, we detected very little evidence either for or against the presence of conjunction memory, and linear trends across ROIs in conjunction memory for Fribbles were either borderline (for the conjunction memory BOLD contrast) or not significant (for the brain-behavior correlation). Thus, although data patterns in Fribbles were numerically in the predicted directions, our experiment appears to have been underpowered for this stimulus set. Third, in the Scenes dataset, we found substantial evidence for above-zero conjunction memory only in PRC. While this aligns with the prediction that conjunction memory signals should reside in MTL rather than visual cortex, the prior literature dissecting specific MTL contributions implicates HC and PHC in scene processing, and PRC in object processing (e.g., Lee et al. 2005; Lee et al. 2008; Staresina et al. 2013). It is unclear why the most robust conjunction memory for scenes was found in PRC and not HC. One possibility is that the experiment was slightly underpowered even for scenes (e.g., we found evidence neither for *nor against* the presence of conjunction memory in HC), another factor may have been imprecise ROI definitions from the probabilistic atlas, and finally it is possible that PRC in fact plays a role in representing conjunctions for scenes (see, e.g., Buffalo et al. 2006; Hannula et al. 2013; Ross et al. 2018 for engagement of PRC by scenes). Interestingly, although the group mean conjunction memory signal in HC did not exceed zero, the strongest brain-behavior correlation for scene conjunction memory was found in HC (BF of 49.42; see Table 4). So, although a hippocampal conjunction memory signal was not present in all subjects, where it was present it was very strongly to related conjunction memory discrimination.

Finally, we discuss two unusual aspects of the experimental design. First, we used three presentations of each item in the test phase (48 trials for each of Novel, Recombination and Old). This may have led to test-based interference: the familiarity differential that can solve the task (Old > Recombination > New) is greatest immediately after study but diminishes as test presentations are repeated. To investigate test-based interference, we examined whether performance changed from the 1^st^ to the 3^rd^ test presentation, finding no overall change in *d’* in Fribbles, and a slight increase in *d’* in Scenes (in contrast to the decrease expected from interference). Thus, test-based interference was minimal, either because the familiarity differential was not impacted by repeated test presentations (even on the 3^rd^ presentation, Old items have been seen many more times than New items), or because interference was offset by non-mnemonic task-practice effects, or by relying on source memory (e.g., for a Novel item: “this item was seen recently in the test phase, but not during the study phase”). Importantly, however, reliance on source memory should only dampen the predicted neural effects: if responses to all items begin to invoke source memory, all theories of memory would predict hippocampal engagement for all items, which should reduce the BOLD contrasts for both feature and conjunction memory and attenuate any gradient of memory signals across the brain. The presence of detectable gradients, in opposite directions, suggests that participants did not rely upon source memory alone.

A second unusual aspect was that, because Recombination stimuli combined features from different families, participants could exploit the familiarity/novelty of 2-feature conjunctions to identify Old or Recombination items. Furthermore, because each 2-feature conjunction appeared in two different items at study, one might argue that they are acquired through “statistical learning” (in contrast to individual items, which resemble episodic “instances”). However, both 3-feature conjunctions (individual items) and 2-feature conjunctions are shown many times at study, and both constitute a complex representation relative to individual features. Although repetition is a hallmark of statistical rather than episodic learning, the strongest episodic memories are those that have been rehearsed or repeated (Mayes and Roberts 2001; Bird et al. 2015), e.g., by prior recounting, internal rehearsal, or repeated viewing of a photograph – a process that surely renders them independent of much of their original context. Thus, to distinguish “episodic” from “statistically learned” memories, we can rely upon neither the number of repetitions, nor the degree of original context retained. Here, we eschew the difficult task of defining these classes of memory: rather than claiming to have created episodic instances, we claim only to have created long-term (i.e., far exceeding 30 seconds), declarative memory.

To conclude, the present findings suggest that the neural substrates of long-term, declarative memory may lie outside of the MTL when the to-be-remembered content is sufficiently simple and purely visual. This finding concurs with other recent studies examining long-term visual memory for simple, visual stimuli (López-Aranda et al. 2009; Thakral et al. 2013; Gavornik and Bear 2014; Cooke et al. 2015; Karanian and Slotnick 2018; Ross et al. 2018). Further, we have demonstrated that recognition memory signals can be made to shift location within the brain by systematically manipulating the complexity of the representations needed for recognition. Together, these results erode the “multiple memory systems” notion of a one-to-one mapping between distinct memory systems and distinct neural substrates. Instead, they support a representational account of cognition, in which regions of the ventral visual-MTL pathway do not specialize in circumscribed cognitive functions but contribute flexibly to cognition whenever their representations are important for solving the task.

## Acknowledgments

This work was supported by the National Science Foundation (award #1554871 to RAC). We thank Dave Huber and Jeff Starns for helpful discussions.

## Notes

Conflict of interest: The authors declare no competing financial interests.

### Competing Interest Statement

The authors have declared no competing interest.

### Summary of Updates

Kept results for stimulus sets separate; updated results without removing outliers; reordered list of ROIs for linear trend analyses; updated an author's affiliation/contact information; manuscript text clarified; reorganized text/figures included in manuscript and supplemental files; additional figures for clarity of design.

