## Supplementary Materials for "The locus of recognition memory signals in human cortex depends on the complexity of the memory representations"

**fMRIPrep Preprocessing**

Preprocessing was performed using fMRIPrep 1.5.2 (Esteban et al., 2019; Esteban, Markiewicz, DuPre, et al., 2018), which is based on Nipype 1.3.1 (Esteban, Markiewicz, Johnson, et al., 2018; Gorgolewski et al., 2011). T1w images were corrected for intensity non-uniformity using N4BiasFieldCorrection (Tustison et al., 2010), distributed with ANTs 2.2.0 (Avants et al., 2008). T1w images were then skull-stripped with antsBrainExtraction.sh, using OASIS30ANTs as the target template. Brain tissue segmentation of cerebrospinal fluid, white-matter and gray-matter was performed on the brain-extracted T1w images using FAST (FSL 5.0.9; Zhang et al., 2001). An unbiased registration of the two T1w images using mri_robust_template (FreeSurfer 6.0.1; Reuter et al., 2010) produced a single T1w-reference map used throughout the workflow. Brain surfaces were reconstructed using recon-all (FreeSurfer 6.0.1; Dale et al., 1999) and the brain mask estimated previously was refined with a custom variation of the method to reconcile ANTs- and FreeSurfer-derived segmentations of the cortical gray-matter of Mindboggle (Klein et al., 2017). Spatial normalization to the ICBM 152 Nonlinear Asymmetrical template version 2009c (MNI152NLin2009cAsym; Fonov et al., 2009) was performed through nonlinear registration with antsRegistration (ANTs 2.2.0), using brain-extracted versions of both the T1w reference and the standard space template.

For each of the 12 functional (BOLD) scans per participant (i.e., 12 test runs), a BOLD reference image and its skull-stripped version were generated using a custom methodology of fMRIPrep. Using this BOLD reference, head-motion parameters were estimated using mcflirt (FSL 5.0.9; Jenkinson et al., 2002) and susceptibility distortions were corrected using a custom workflow of fMRIPrep that estimated a deformation field based on a field map that was co-registered to the BOLD reference. This estimated susceptibility distortion was used to calculate an unwarped BOLD reference for more accurate co-registration with the T1w reference using bbregister (FreeSurfer). In a single interpolation step that concatenated the above transformations, the BOLD time-series were resampled into subject native space. Volumetric (gridded) resamplings were performed using antsApplyTransforms (ANTs), configured with Lanczos interpolation to minimize the smoothing effects of other kernels (Lanczos, 1964). Surface (nongridded) resamplings to subject native space (fsnative) and the fsaverage template space were performed using mri_vol2surf (FreeSurfer) and the resulting transformation matrices were later used to apply probabilistic ROI atlases. For more details, see the workflows section in fMRIPrep’s documentation.

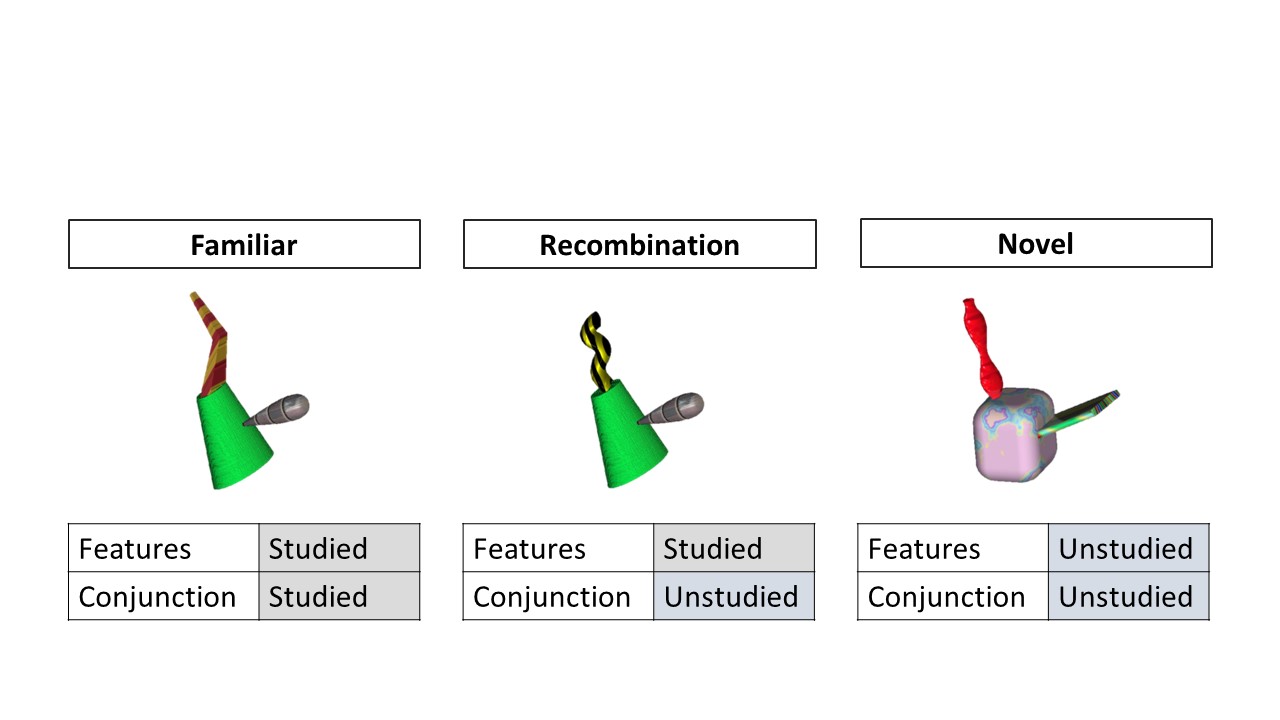

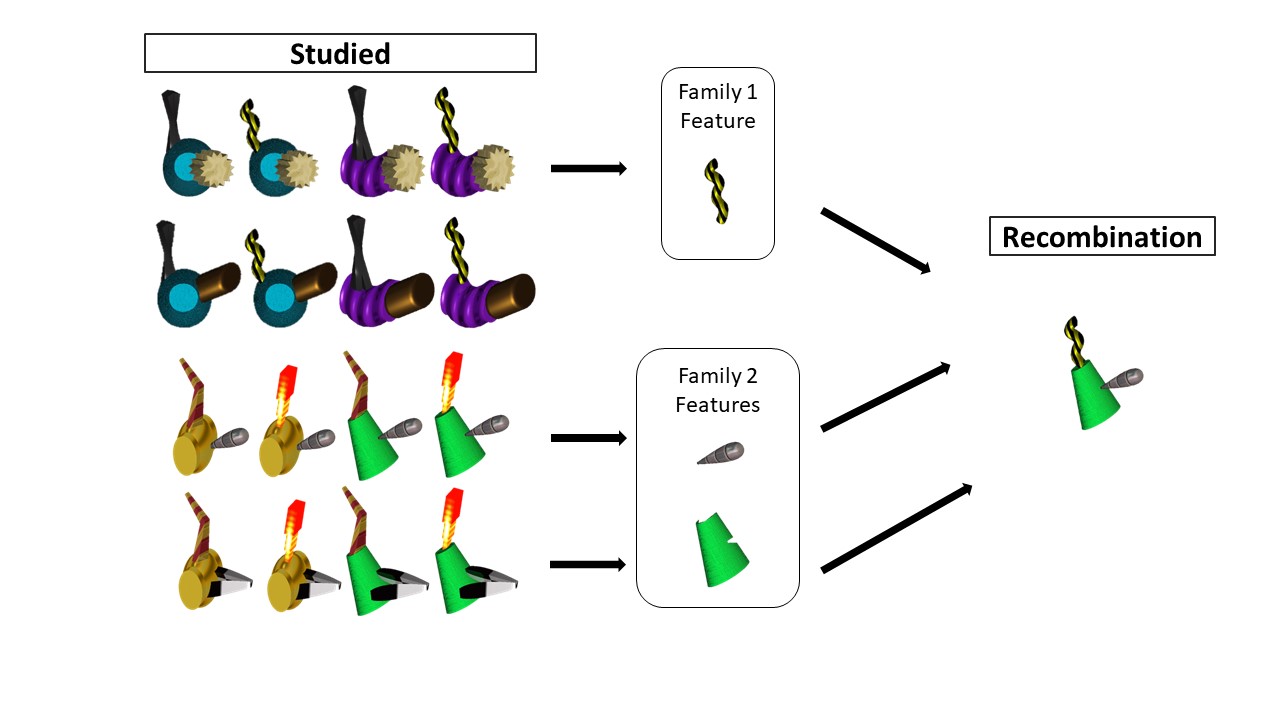

**Figure S1. Fribble Stimulus Set Recombination Examples.** *Top*: the three mnemonic stimulus classes differed according to whether features and the conjunction of features were studied in the study phase; *Bottom*: counterbalanced across participants, two of the Fribble families were designated to be studied (i.e., presented in the study phase). In this example, a Recombination stimulus is created by combining the tail feature from the first family with the body and head features from the second family. All potential 1 + 2 feature combinations within a given pair of families yielded 48 Recombination stimuli.

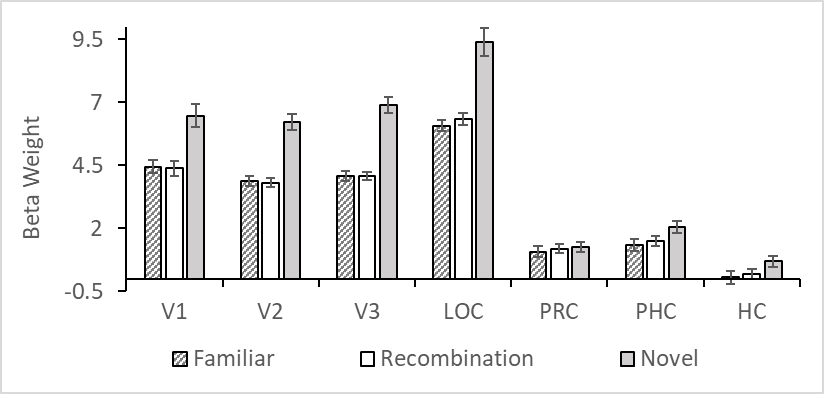

**Figure S2. Mean Beta Weights for Fribbles.** Beta weights were used in the calculation of memory scores. Specifically, for Feature Memory, beta weight estimates from the GLM for correct Recombination trials were subtracted from beta weight estimates for correct Novel trials. Similarly, for Conjunction Memory, beta weight estimates for correct Familiar trials were subtracted from beta weight estimates for correct Recombination trials. Error bars are within-subject SEM.

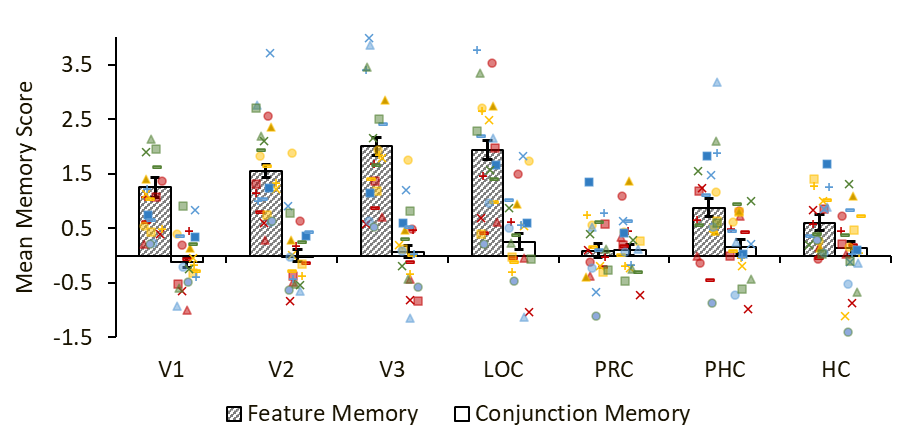

**Figure S3. Memory Scores with Individual Participant Points for Fribbles.** Memory scores, as measured by Cohen’s *d*, for the contrast of Novel and Recombination trials (i.e., Feature Memory) and the contrast of Recombination and Familiar trials (i.e., Conjunction Memory). Striped and solid white bars show group means; points show individual participants, where each unique color-marker combination depicts the same individual across ROIs. Error bars are within-subjects SEM.

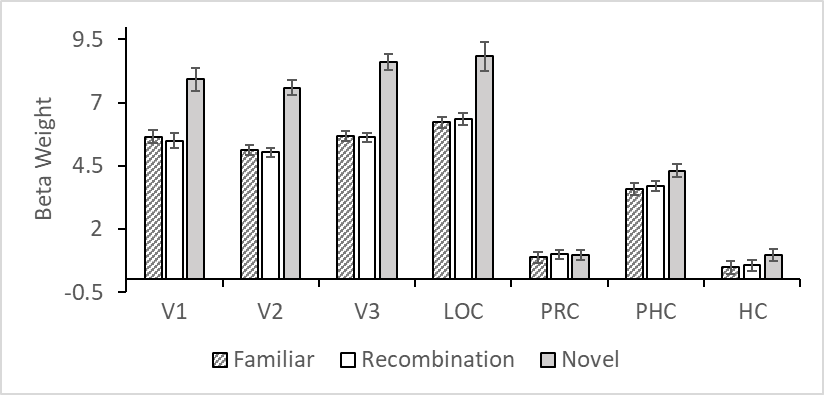

**Figure S4. Mean Beta Weights for Scenes.** Beta weights were used in the calculation of memory scores. Specifically, for Feature Memory, beta weight estimates from the GLM for correct Recombination trials were subtracted from beta weight estimates for correct Novel trials. Similarly, for Conjunction Memory, beta weight estimates for correct Familiar trials were subtracted from beta weight estimates for correct Recombination trials. Error bars are within-subject SEM.

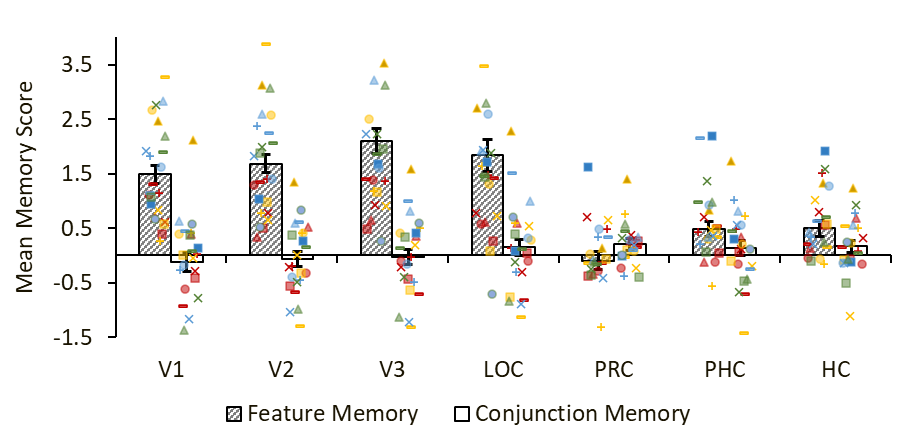

**Figure S5. Memory Scores with Individual Participant Points for Scenes.** Memory scores, as measured by Cohen’s *d*, for the contrast of Novel and Recombination trials (i.e., Feature Memory) and the contrast of Recombination and Familiar trials (i.e., Conjunction Memory). Striped and solid white bars show group means; points show individual participants, where each unique color-marker combination depicts the same individual across ROIs. Error bars are within-subjects SEM.
